# A Stochastic Population Model for the Impact of Cancer Cell Dormancy on Therapy Success

**DOI:** 10.1101/2023.12.15.571717

**Authors:** Jochen Blath, Anna Kraut, Tobias Paul, András Tóbiás

## Abstract

Therapy evasion – and subsequent disease progression – is a major challenge in current oncology. An important role in this context seems to be played by various forms of cancer cell dormancy. For example, therapy-induced dormancy, over short timescales, can create serious obstacles to aggressive treatment approaches such as chemotherapy, and long-term dormancy may lead to relapses and metastases even many years after an initially successful treatment. The underlying dormancy-related mechanisms are complex and highly diverse, so that the analysis even of basic patterns of the population-level consequences of dormancy requires abstraction and idealization, as well as the identification of the relevant specific scenarios.

In this paper, we focus on a situation in which individual cancer cells may switch into and out of a dormant state both spontaneously as well as in response to treatment, and over relatively short time-spans. We introduce a mathematical ‘toy model’, based on stochastic agent-based interactions, for the dynamics of cancer cell populations involving individual short-term dormancy, and allow for a range of (multi-drug) therapy protocols. Our analysis shows that in our idealized model, even a small initial population of dormant cells can lead to therapy failure under classical (and in the absence of dormancy successful) single-drug treatments. We further investigate the effectiveness of several multidrug regimes (manipulating dormant cancer cells in specific ways) and provide some basic rules for the design of (multi-)drug treatment protocols depending on the types and parameters of dormancy mechanisms present in the population.

## 1. Introduction

### 1.1. Motivation

Cancer is a highly complex disease that is among the leading causes of deaths worldwide. Despite a range of different therapeutic approaches, a major challenge in current oncology seems to be the emergence of cancer therapy evasion, frequently resulting in treatment failure and relapses. In several cases, such therapy resistance has been linked to forms of *cancer cell dormancy* (cf. eg. [F+23] and the references therein), by which we mean a phenotypic switch into a non-proliferating and protected (therapy-evading) cellular state of reduced activity. Such transitions can be triggered by the stress resulting from chemo-, immuno- and even radiation therapy, but may also happen spontaneously or as a result of other environmental cues [DRS21]. The emergence of dormant sub-populations over relatively short timescales can lead to therapy resistance, while long-term dormancy may result in relapses and metastases many years after the initial cancer formation [AG07, PC20]. In this context, the somewhat unclear boundaries between different dormancy-related mechanisms and concepts, including quiescence, persistence, latency, tolerance and stemness, seem to further complicate the derivation of general strategies to handle the resulting therapeutic challenges (see e.g. [V19, F+23] for comments on the related nomenclature).

Due to their apparently significant role in therapy failure, dormancy mechanisms have recently attracted increasing interest in the literature, where a wide range of mechanisms and strategies have been described (see e.g. [AG07, F+23, RM19, SD21, V19, EI19, EL15]). Beyond cancer biology, dormancy-related survival and coexistence strategies are currently also being studied in many other contexts, including mathematical modelling and studies in population genetics, evolution, and ecology, see e.g. [LdHWB21] for an overview. There, it becomes increasingly clear that the population-level effects of dormancy are heavily-influenced by the particular type of (abstract) switching mechanism, including in particular spontaneous, responsive and competition-induced dormancy transitions, see e.g. [SL18, BT20, BHS21] for some general results in this direction.

In the present working paper, our starting point is to incorporate similarly paradigmatic dormancy mechanisms into a (highly idealized) individual-based stochastic cancer population model, while augmenting its dynamics with different (multi-drug) agent-based treatment regimes. We then investigate the consequences of these micro-interactions for the limiting dynamics of the macroscopic *population-level model*, in particular regarding the emergence of therapy failure over short time-scales, and the relative effectiveness of different treatment protocols – all in basic scenarios.

Ultimately, we hope to indicate a way to gain systematic insights into the population-level effects of dormancy that may help to optimize treatment protocols (in more realistic scenarios) in order to tackle some of the challenges posed by dormancy-related therapy resistance. One concrete way to achieve this could be the design of combination therapies where multiple drugs will be administered in a way that is tailored to the underlying dormancy mechanisms and resulting population dynamics while at the same time obeying constraints resulting e.g. from negative side-effects.

### 1.2. Modelling approach, and basic transitions related to dormancy and treatment

Our starting point will be a multi-type stochastic *birth-death process with interactions*, describing the different types of cancer cell populations (including dormant and actively proliferating states) as well as multiple drug agents administered in externally controlled treatment regimes. This model will be *mechanistic* and incorporate in particular various forms of abstract dormancy strategies. The transitions will differ e.g. regarding the birth, death and switching rates, the drug sensitivities and supply rates, the option and rate of spontaneous dormancy emergence, and the interplay with other cells (competition) and drugs (therapy-induced switches). Deriving an explicit class of models thus immediately necessitates some basic classification and formalisation of dormancy mechanisms.

A first abstract classification can be based on the distinction between *individual cell dormancy* and *tumour mass dormancy*. The latter describes the tumour *population level* equilibrium of reproduction and apoptosis (see e.g. [F+23, AG07, EI19, EL15] and the references therein), and is not the focus of this paper. Instead, we consider individual cell dormancy, where *single cancer cells* switch into a non-proliferating and protected state of reduced activity with certain rates according to mechanisms that we will specify below.

A second classification can be based on the involved time-scales of dormancy. Here, we focus on *short-term dormancy* (e.g. given by cell-cycle arrest or phenotypic switches into quiescence) that can be modelled by transition rates that are of the *same order* as the other interactions of our system. Hence, we do not consider long-term dormancy in the form of “metastatic latency”, such as dormant disseminated tumour cells that rest in certain niches for many years until changes in their local environment lead to their resuscitation.

Finally, to incorporate short-term cell dormancy into our model, we need to abstractly describe the related *switching mechanisms*. The distinction between the following transitions will be crucial for the macroscopic behaviour of the system.

Regarding the *initiation of dormancy*, one may distinguish

- spontaneous (stochastic) switching into dormancy, which may lead for example to the presence of dormant / resistant sub-populations even in therapy-na ï ve tumours,
- dormancy triggered in response to chemo-, immuno- or radiation therapy,
- dormancy initiation due to internal competition / overcrowding / starvation,
- and dormancy initiation due to further environmental cues.

Regarding the *maintenance of dormancy* or *re-activation*, one may distinguish between

- spontaneous (stochastic) resuscitation,
- drug-induced resuscitation,
- drug-delayed resuscitation, e.g. due to quiescence-inducing drugs,
- resuscitation due to external environmental cues or population size fluctuations.

In our model, we will allow most of the above transitions, but will refrain from introducing responsive switches due to external environmental cues other than drug therapy.

While these distinctions in relation to dormancy mechanisms already indicate the complexity of the field, one could easily incorporate more features such as further cell phenotypes, age- and spatial structure, stem-like behaviour, or the emergence and accumulation of resistance mutations. However, since our focus for this paper is on the basic effects of dormancy mechanisms, we refrain from incorporating these additional ramifications.

Regarding possible *treatment regimes*, we will incorporate time-dependent external drug administration on the individual level (“agent-based”) of our model, allowing for multiple (different) drugs each with specific impact on the active and dormant cancer cells.

Basic *therapeutic targets* can be specified as the *remission of the tumour*, in the sense that the total number of tumour cells decreases below a certain threshold in a given period of time, or by *limiting the number of resistance mutations* that can be accumulated by the cancer cell population, again within a certain period of time. Note that the idea behind the second goal is to reduce the probability of the emergence of a resistance mutation leading e.g. to the emergence of aggressively proliferating therapy-resistant cancer cells (TRCC). It will turn out that, depending on the details of the dormancy-related mechanisms, the two specific goals may necessitate different treatment protocols, in particular since the mutation rate in the dormant state may be lower, but (somewhat counter-intuitively) also be higher than in the active state [R+22].

### 1.3. Organisation of the paper

In Section 2, we formally introduce the stochastic individual-based mechanisms of our interacting birth-death process, including the externally determined treatment protocols. We then derive the corresponding many-particle limits of our model, leading to a population-level dynamical system, based on a suitable convergence theorem. For the stochastic system without treatment, we also investigate analytically the probability of extinction in the early phase of the cancer emergence, and the rate of growth in the case of a small tumour.

In Section 3 we investigate, mostly by simulation, how dormancy traits can lead to therapy evasion, and, given the details of the dormancy strategies, how to optimize treatment protocols potentially involving multiple drugs. Optimization criteria can e.g. be the minimization of the probability of the emergence of a resistant type before remission under certain conditions on side-effects. Besides the specific results, we aim to formulate *general observations* (GOs) for the impact of cancer cell dormancy on therapy success in our highly idealized scenario.

The paper concludes with a discussion (Section 4), in which we review our results, the limits of our model, the choice of parameters, possible extensions, and relations to other work.

## 2. A Population Model For Cancer Cell Dormancy And Multi-Drug Treatment

We begin with a stochastic model for the basic interactions between active and dormant cancer cells, and the agents of two drugs acting in specific ways on these active and dormant cells. Extensions to additional cell types and multiple drugs are in principle straightforward.

### 2.1. The basic interacting birth-death process and its scaling limit

Our stochastic model is given by a continuous-time Markov chain *{N* (*t*)*}*_*t≥*0_ on 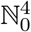of the form

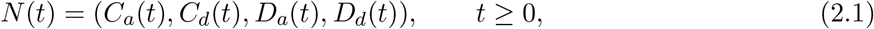

where the quantities on the right-hand side describe, at time *t ≥* 0, the number of active cancer cells *C*_*a*_(*t*), of dormant cancer cells *C*_*d*_(*t*), of agents of a drug acting on active cancer cells *D*_*a*_(*t*), and of agents of a drug acting on dormant cancer cells *D*_*d*_(*t*). To facilitate a subsequent many particle-limit, let *K >* 0 be a parameter denoting the overall *carrying capacity* of our population. We allow for spontaneous/stochastic switching between active and dormant cells as well as drug-induced responsive switching. The individual dynamics and parameters of our systems are then as follows (visualized also in Figure 1(a)–(f)).

**Figure 1.**
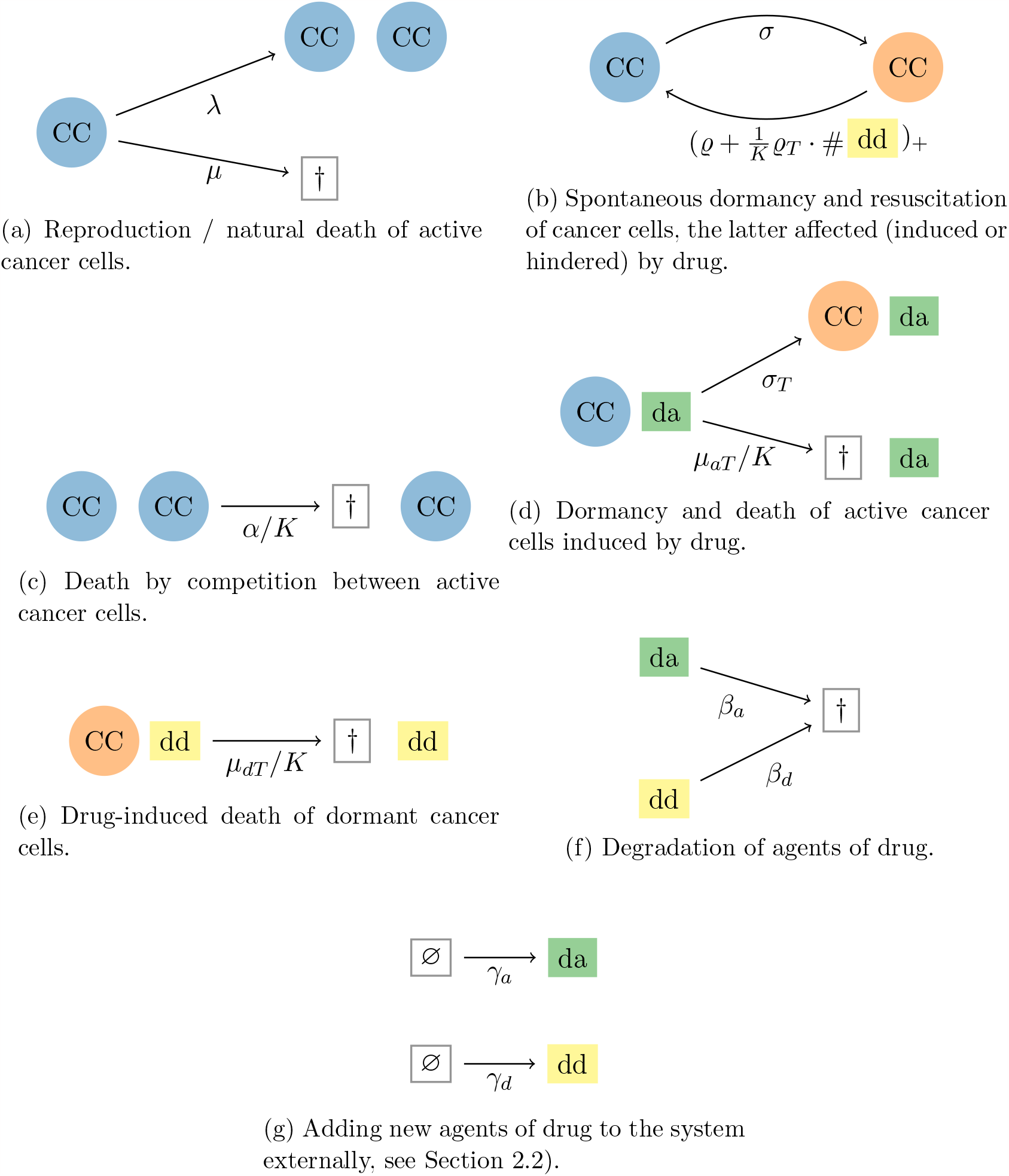
Graphical representation of the transitions of the stochastic model from Section 2.1, involving transitions (a) to (f), and Section 2.2, additionally involving (g). Circles denoted by CC represent cancer cells (blue: active state; orange: dormant state), while rectangles represent the agents of the drug targeting active (da) and dormant (dd) cells, respectively. The symbol *†* indicates cell death or drug degradation. The symbol ∅ represents an external source for the agent of a drug.

#### Transitions of active cancer cells

- An active cancer cell undergoes cell division into two active cancer cells at rate *λ >* 0.
- An active cancer cell dies naturally at rate *μ*, where 0 *< μ < λ*.
- An active cancer cell dies due to drug treatment at rate *μ*_*aT*_ *D*_*a*_*/K*, where *μ*_*aT*_ *>* 0 describes the effectiveness of the drug.
- An active cancer cell dies due to competition with other active cancer cells at rate *αC*_*a*_*/K* with *α ≥* 0.
- An active cancer cell switches into dormancy spontaneously at rate *σ >* 0.
- An active cancer cell (optionally) switches responsively into dormancy due to drug treatment at rate *σ*_*T*_ *D*_*a*_*/K* with *σ*_*T*_ *≥* 0.

#### Transitions of dormant cancer cells

- A dormant cancer cell spontaneously switches back (“resuscitates”) into an active cancer cell at rate *ϱ* with *ϱ >* 0.
- A dormant cancer cell responsively switches back into an active cancer cell due to resuscitation promoted by drug treatment at rate *ϱ*_*T*_ *D*_*d*_*/K* with *ϱ*_*T*_ *≥* 0.
- If the drug targeting dormant cancer cells suppresses instead of promotes resuscitation, the above rates will be replaced by the combined rate (*ϱ* + *ϱ*_*T*_ *D*_*d*_*/K*)_+_ with *ϱ >* 0, *ϱ*_*T*_ *≤* 0.
- A dormant cancer cell dies due to treatment (optionally) at rate *μ*_*dT*_ *D*_*d*_*/K* with *μ*_*dT*_ *≥* 0.

#### Dynamics of agents of drugs

- An agent of the drug targeting active cancer cells degrades at rate *β*_*a*_ *>* 0.
- An agent of the drug targeting dormant cancer cells degrades at rate *β*_*d*_ *>* 0.

Note that the rates *σ*_*T*_, *ϱ*_*T*_ and *μ*_*dT*_ may be equal to zero, which allows for models where the respective effects of the drugs are absent. Further, if *ϱ*_*T*_ is strictly negative, which corresponds to the scenario when the drug suppresses resuscitation, then the overall resuscitation rate of dormant cancer cells may equal 0, but of course cannot become negative.

At the moment, we have not yet specified how the external drug administration occurs, so that the initially present number of agents *D*_*a*_(0) and *D*_*d*_(0) will simply degrade by following pure-death processes over time. In the next subsection 2.2 we will present variants of the model with external drug administration.

To obtain the *many-particle limit* of our system, which corresponds to letting *K → ∞*, we may employ standard arguments (see e.g. [EK86, Theorem 11.2.1]). Indeed, the vector of rescaled population sizes

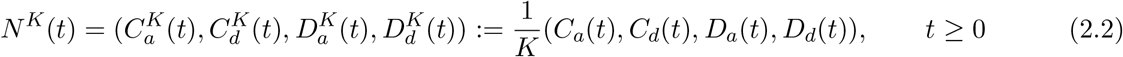

converges a.s. uniformly on compact time intervals of the form [0, *T*] to the solution *n*(*t*) = (*c*_*a*_(*t*), *c*_*d*_(*t*), *d*_*a*_(*t*), *d*_*d*_(*t*)) of the system of ODEs

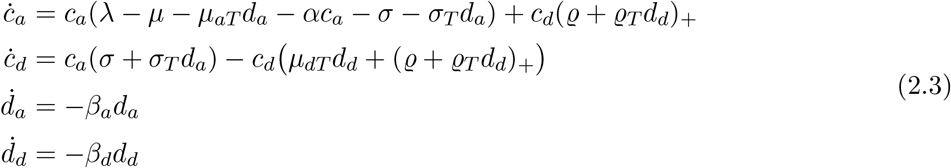

under the condition that we have the convergence of initial conditions *N* ^*K*^(0) *→ n*(0) a.s. as *K → ∞*. This system resembles a classical logistic growth model with an additional dormancy compartment.

### 2.2. Incorporating treatment protocols

Our treatment modelling is motivated by chemotherapy, which is typically a “pulsed” treatment, that is, it is applied periodically. We thus additionally incorporate periodic increases of the functions *d*_*a*_, *d*_*d*_ and interpret them as external drug administration. Two special cases will be particularly important.

a. *Instantaneous drug administration*. Suppose that large quantities of agents of a drug are administered instantaneously (e.g. via “injection”) at periodic time-points with interval length *ω*. If the number of administered agents, say at time 0, is of size *Kd*_*a*,0_ *>* 0 (resp. *Kd*_*d*,0_ *>* 0), this will lead to jumps of size 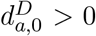 (resp. 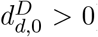) in the limiting system. Immediately after these jumps, the administered chemicals decay over time with rate *β*_*a*_ (resp. *β*_*d*_) according to the previous dynamics, so that the shape of the drug concentration after the first jump (here at time 0) takes the form

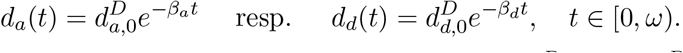

Further instantaneous applications leading again to jumps of size 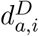(resp. 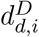) at the *i*-th multiple of *ω* will accumulate accordingly.
b. *Continuous drug administration*. In contrast, if the drug is administered continuously (e.g. via “infusion”) over an extended time interval, we add an external “immigration component” to the system, corresponding to Figure 1 (g):
  - At rate *γ*_*a*_(*t*) *≥* 0, new agents of the drug targeting active cancer cells enter into the system. Here, in contrast to all previous rates, we allow for externally determined time-dependent immigration rates.
  - At rate *γ*_*d*_(*t*) *≥* 0, new agents of the drug targeting dormant cancer cells immigrate into the system. Again, the immigration rate is externally determined and depends on time.

In this case, we obtain a slightly different scaling limit, namely

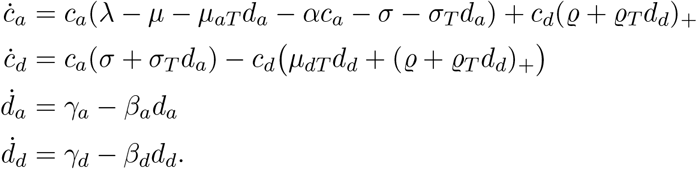

One possible scenario is to apply an infusion from time zero up to some time *S*_*a*_ *>* 0 in the case of the drug targeting active cancer cells, and *S*_*d*_ *>* 0 in the case of the drug targeting dormant cells. Then, we obtain for 0 *≤ t ≤ ω* that

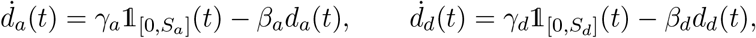

where *γ*_*a*_, *γ*_*d*_ *>* 0 are constants determining the rate of inflow of the drugs. This is an inhomogeneous linear ODE of first order in one dimension. Using the variation of constants formula and taking into account the initial condition *d*_*a*_(0) = 0, we obtain

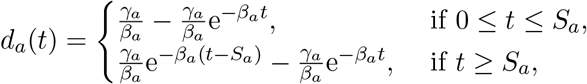

and analogously for *d*_*d*_(*t*). In the case of multiple drug infusions within (disjoint) time intervals, we replace 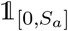 resp. 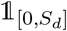 with the sum of the indicators of the corresponding intervals. This is e.g. the case if treatment is repeated with some period *ω ≥ S*_*a*_ resp. *ω ≥ S*_*d*_.

### 2.3. Convergence to a limiting dynamical system under externally specified periodic treatment

We consider the set-up of Section 2.2 and show that the corresponding Markov chain, now with additional (externally controlled) instantaneous drug administrations (“injections”), converges to a (suitably modified) dynamical system with corresponding jumps in drug concentrations.

To this end, assume that treatment will be administered at times 0 = *T*_0_ *< T*_1_ *< · · ·*, and that the instantaneously administered doses for the drug targeting active (respectively dormant) cancer cells are of the from 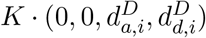 for some 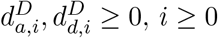. To obtain the desired limiting system, we employ the classical convergence result from [EK86] in a piecewise fashion on the intervals between consecutive drug administrations.

#### Lemma 2.1

*Let T*_*i*_, *i ≥* 0, *be the times of instantaneous drug administrations. Assume that the convergences of the initial conditions*

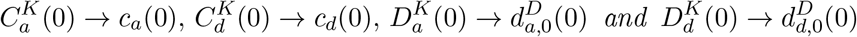

*hold almost surely as K → ∞. Let* 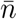 *be the unique piecewise continuously differentiable function such that*

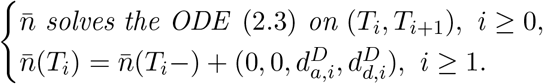

*Then, for all T ≥* 0, 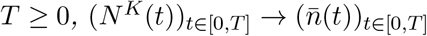 *uniformly almost surely, i.e*.

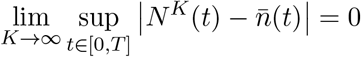

*almost surely*.

The simple proof of this lemma can be found in Section 5.1 in the Appendix. A corresponding result applies to the case of continuous “infusions” corresponding to immigration rates *γ*_*a*_, *γ*_*d*_ *≥* 0 during time intervals of the form [*T*_*i*_, *T*_*i*_ + *S*_*a,i*_], [*T*_*i*_, *T*_*i*_ + *S*_*d,i*_] contained in [*T*_*i*_, *T*_*i*+1_]. In that case, an explicit solution for the drug concentrations *d*_*a*_, *d*_*d*_ (which do not depend on *c*_*a*_, *c*_*d*_) can be obtained via the variation of constants formula.

Note that the *T*_*i*_’s can also be chosen as random finite stopping times (uniformly in *K*).

### 2.4. Probability of extinction and rate of growth under spontaneous switching

Interesting quantities that can be evaluated analytically are the probability of extinction and the rate of growth in the early phase of cancer/tumour emergence, that is, when the cancer cell population is still small enough such that competitive effects are negligible and can thus be described by a super-critical branching process. For concreteness, we denote this two-type branching process by 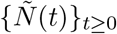. It takes values in 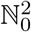 and is of the form

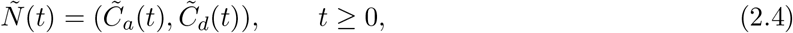

where 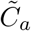 and 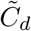only undergo the dynamics corresponding to transitions (a) and (b) in Figure 1.

Now, recall from elementary theory of multi-type branching processes (see eg. [AN72, Chapter V.3]) that the *probability of extinction* of the cancer cell population when starting from a single active cell – denoted by 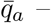 and when starting from a single dormant cell – denoted by 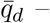 is given by the solution (*q*_*a*_, *q*_*d*_) in [0, 1]^2^ *\ {*(1, 1)*}* of the system

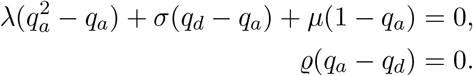

We can easily solve this to find

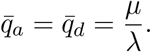

This is *exactly* the probability of survival that we see for a population without dormancy. The result comes as no surprise because the dormancy mechanism prevents reproduction and hence only freezes the individuals. Hence, it may take longer for a population of cells to go extinct, but the probability of it happening remains the same.

Concerning the initial *exponential rate of growth*, this quantity is determined by the principal eigen-value *λ*_1_ of the offspring mean matrix

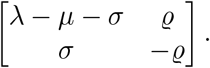

Gershgorin’s circle theorem then implies that the principal eigenvalue is less than *λ − μ* (for *λ > μ*). In particular, spontaneous dormancy indeed decreases the rate of growth of a tumour. To get a quantitative intuition about the strength of this effect, assume that we increase the rates of dormancy and resuscitation simultaneously, now considering *ρx* and *σx* for *x → ∞*. This corresponds to the mean matrix

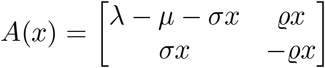

with eigenvalues *λ*_1_(*x*) *> λ*_2_(*x*) given by

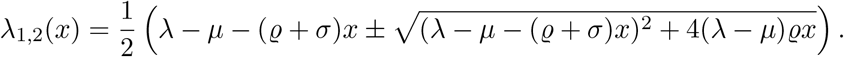

Using *λ*_1_(*x*) = det(*A*(*x*))*/λ*_2_(*x*), it is easy to show that

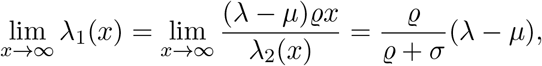

where we used that *λ*_2_(*x*) *∼ −*(*ϱ*+*σ*)*x* as *x → ∞* in the second equality. This is to say that as switching becomes infinitely fast, the growth rate *λ*_1_(*x*) is lowered from *λ*_1_(0) = *λ − μ* (which corresponds to the case without dormancy) to a fraction of this rate, where the coefficient *ϱ/*(*ϱ* + *σ*) is precisely the value of the active component of the stationary distribution of a non-reproducing system that merely switches between active and dormant states with rates *ϱ* and *σ*.

Further, the dependence on *x* is monotone, i.e.

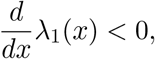

and we also have lim_*x↓*0_ *λ*_1_(*x*) = *λ − μ*. This means that the presence of dormancy does not have a strong impact for small *x* and only becomes significant once the switching happens sufficiently fast compared to the reproductive rate. Indeed, when active cells replicate, the two resulting cells are both initially active. Hence, when the rate at which an individual cell enters a phase of dormancy is low, this very same cell undergoes almost as many cell divisions as it would without the possibility of becoming dormant.

Our system (2.3) with drug administrations has a coefficient matrix on the right-hand side similar to *A*(*x*) if we solve the independent equations for drug concentrations and neglect the competition term. However, this system then becomes non-autonomous, and in general cannot be solved analytically.

Instead, we will perform a simulation study to investigate the effects of dormancy on treatment protocols in the next section.

## 3. Consequences of dormancy for therapeutic approaches: challenges and results

### 3.1. Evaluating treatment success

From equation (2.3) we already know the dynamics of the cancer cell population / tumour size on each time interval between consecutive drug administrations. To evaluate the effectiveness of treatments, we further need to specify their constraints and precise therapeutic goals, which will then lead to an *optimal control problem*. Two natural objectives are the minimization of

1. the total cancer cell population / tumour size by the end of the treatment, and
2. the probability of the occurrence of a resistance mutation.

The *final tumour size* is quite an obvious criterion for treatment efficiency and can easily be translated e.g. into the objective function

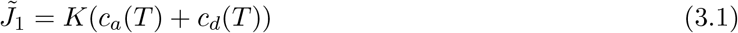

for some *T >* 0 determining the time at which the treatment ends. However, when considering periodic treatment protocols with different interval lengths *ω*, 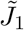 may not be a suitable choice. For example, some treatments may have their last dose administered right before *T*, while others have them significantly before *T*, which may lead to different final cancer cell numbers at *T* and imply an “unfair” comparison of these treatments. To account for such errors, we will instead consider the *average number of cells* at the end of treatment when averaging over the last 30 days. More precisely, this leads us to the objective function

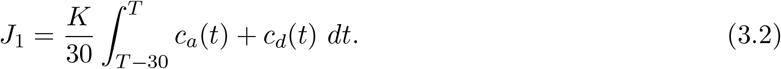

Of course, other choices are also well possible.

Concerning the *probability of the occurrence of a resistance mutation*, we assume that throughout treatment, each cancer cell has independently of all other cells a small probability for mutating into a treatment resistant type, say *m*_*a*_ *≥* 0 per day for active cells, and *m*_*d*_ *≥* 0 per day for dormant cells. In order to minimize the probability for the emergence of a resistance mutation, one may then aim to minimize the objective function

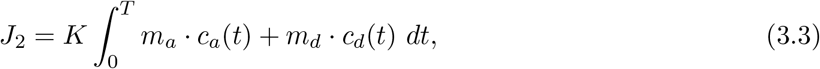

which is proportional to the expected total number of mutations accumulated up to time *T*. The two main scenarios for the occurrence of mutations are errors during DNA repair as well as DNA replication, and we will assume both of these mutation mechanisms to be captured by the mutation probabilities *m*_*a*_ and *m*_*d*_. We will also refer to the above weighted integral of *c*_*a*_ and *c*_*d*_ as *area under the curve* (AUC).

Having specified these prototypical objective functions, we further need to consider constraints for the control functions *d*_*a*_ and *d*_*d*_. Since, we focus on *pulsed periodic treatments* of finite length, these are uniquely determined by their decay rates *β*_*a*_ and *β*_*d*_, the period length *ω*, total treatment length *T* (equivalently, the total number of administrations) and the drug doses 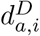and 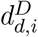. To simplify these constraints even further, we will additionally assume for each drug that the doses given in each treatment cycle are constant, i.e. 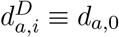 and 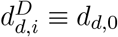. In order to make these doses comparable for different choices of time periods *ω*, we will normalize them and work with *daily doses*. Given a daily dose *d*_*·*,daily_ and a period length *ω*, the effective dose given at the beginning of each treatment cycle can be calculated by

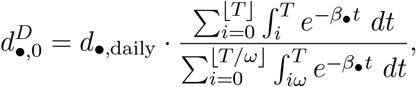

where • is a placeholder for either ‘*a*’ or ‘*d*’. Here, each summand in the sums of both the numerator and denominator is the total contribution of the drug administered at the *i*-th cycle to the total drug in the system administered throughout the treatment with daily administration and administrations at time intervals of length *ω* respectively. Hence, for different periods the daily dose allows us to determine the dosage administered at intervals with period *ω* such that the total drug concentration remains constant. Our constraints will now be

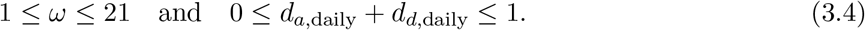

Thus, we consider the optimal control problem consisting of minimizing *J*_1_ or *J*_2_ under the dynamics 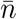 from Lemma 2.1 subject to (3.4). For concreteness, we will say that treatment is successful if *J*_1_ *≤* 100 or *J*_2_ *≤* 1. The rationale behind these choices is that there might be a reasonable chance for a tumour to be eradicated due to stochastic fluctuations once it is of sufficiently small size for a prolonged period of time (here, below 100 cells on average). Similarly, a treatment could be considered to be successful if the expected number of emerging resistance mutations during treatment is sufficiently low, e.g. less than one in our case.

Due to the way that the control functions are designed, unfortunately we cannot make use of standard optimal control theory to tackle our problem analytically. Usually, the control functions will be *L*^*∞*^-functions with the limitations being determined by their extreme values or some other general constraints (e.g. integral constraints) directly related to them. Due to the predetermined shape of our control functions we are operating in a small subset of *L*^*∞*^-functions which we cannot characterize precisely using path constraints. Further, the constraints on the control functions are of an indirect nature since they do not consider the explicit path of the drug concentrations.

### 3.2. Therapy failure due to dormancy

Before discussing how different treatment protocols may deal with dormancy, we illustrate how dormancy may diminish the success of a treatment protocol that is successful in absence of dormancy. For this, we assume from now on the following parameter choices (unless stated otherwise), which are discussed in Section 4:

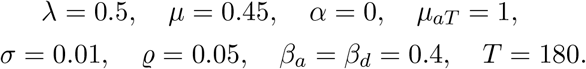

A summary of all parameters (fixed and variable) can be found in Table 1 in the appendix. Furthermore, we will assume the tumour to be treated immediately and the initial size of the tumour to be 10^8^ cells. With a slight abuse of notation, we identify *K · c*_*a*_ and *K · c*_*d*_ from Lemma 2.1 with *c*_*a*_ and *c*_*d*_, respectively. The factor *K* determines the effective population size which we can incorporate directly into the solution of the differential equation (2.3). We can do this because neither *c*_*a*_ nor *c*_*d*_ have a quadratic term in (2.3) due to our choice of *α* = 0 that we will comment on below. In particular, we say that *c*_*a*_(0) + *c*_*d*_(0) = 10^8^. This choice of parameters means that we consider treatment over the period of roughly 6 months for a tumour which has a doubling time of about 14 days. We call cancer cells dormant when they undergo a prolonged period without reproduction. In our case this is reflected by our choice of *ϱ* which in expectation has dormant cells inactive for 20 days. A tumour might typically be composed mostly of active cells, hence we chose *σ* = 0.01 to have a ratio of 1:5 active to dormant tumour cells (i.e. 1*/*6 of the total tumour cells are dormant due to spontaneous switching). The rates for drug decay are such that the half-life of the drugs are roughly 2 days.

**Table 1.**
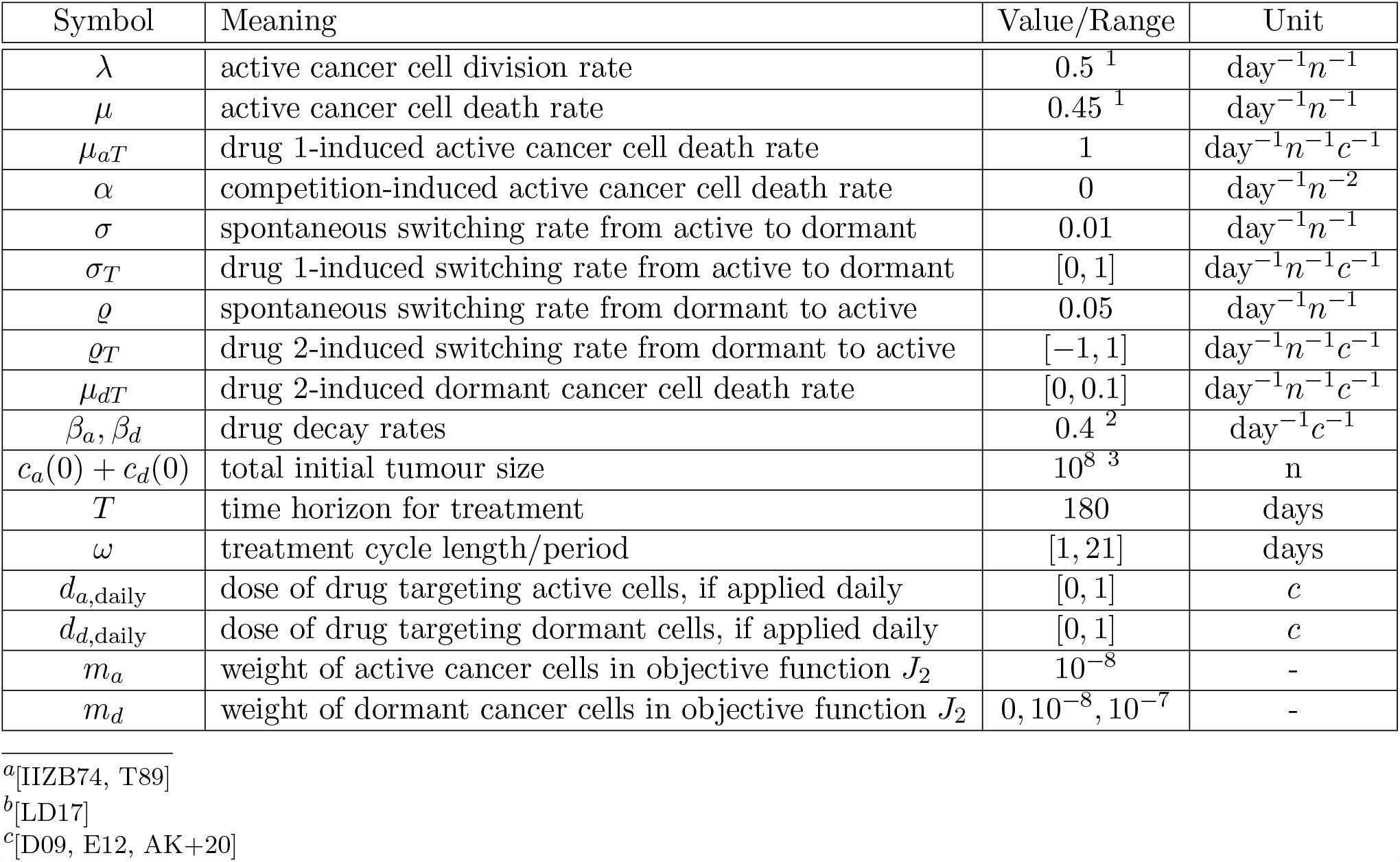
Parameters for our simulation study.

Observe that we assume *α* = 0 which turns off the competition between active cancer cells and would correspond to unbounded exponential growth of the tumour. This can be justified since we are interested in modelling the treatment of a tumour while it is still reasonably small. Because a tumour will commonly reach its maximal size only after it has become lethal, we assume our tumour to be so small initially that the competition term is indeed negligible.

For now, consider a treatment with one drug administered at a daily dose of *d*_*a*,daily_ = 0.1 and a treatment period of *ω* = 14 in the absence of dormancy, i.e. *c*_*a*_(0) = 10^8^, *c*_*d*_(0) = 0 and *σ* = 0. Then the tumour is driven to extinction, i.e. *J*_1_ *<* 1, after the 7th treatment cycle. This is shown in Figure 2 (left).

**Figure 2.**
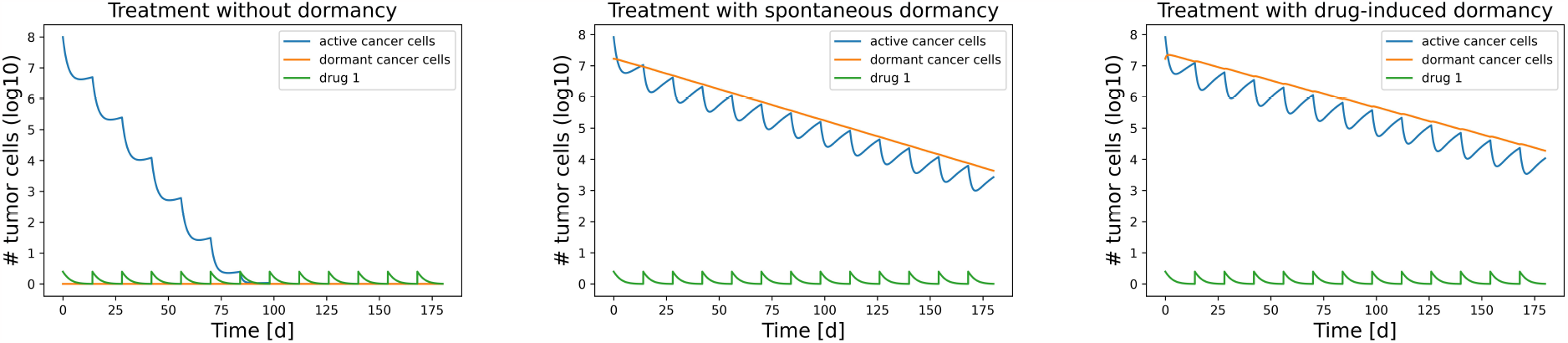
Dynamics of tumour size and drug concentration over time plotted on logarithmic scale. Left: In absence of dormancy, treatment leads to extinction of the tumour. Middle: With spontaneous dormancy, tumour remains of considerable size at end of treatment. Right: With additional drug-induced dormancy, treatment outcome is even worse.

However, the presence of a dormant sub-population may change the outcome of this treatment plan drastically. In order to demonstrate this, we assume the tumour to initially have an equilibrium of active and dormant cells. This can be calculated to be

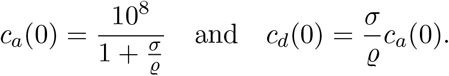

Dormant cells cannot be targeted by the drug any more and hence the tumour cannot be eradicated at a rate faster than the resuscitation rate *ϱ*. This leads to failure of treatment as is illustrated by Figure 2 (middle). The situation gets even worse if active cancer cells may switch into dormancy at an increased rate in response to the stress caused by the drug administration, see e.g. [EL15, R+22]. Indeed, if we assume that *σ*_*T*_ = 0.1, i.e. a drug-induced switching rate that (multiplied by the drug concentration) is higher than the spontaneous switching rate *σ* = 0.01 for a significant time after drug administration, then the final tumour size increases even further. We depict this in Figure 2 (right).

The fact that the presence of dormant cancer cells can prevent treatment success leads us to investigate the maximum equilibrium fraction of dormant cells in a tumour such that a given treatment plan can still be successful.

For concreteness, recall that we consider treatment to be successful if *J*_1_ *≤* 100, i.e. on average there are less than 100 cells over the last 30 days of treatment. Then, for fixed *ϱ* = 0.05 (and given *ω* and *d*_*a*,daily_) we compute the largest *σ* such that *J*_1_ *≤* 100. For these *ϱ* and *σ*, the corresponding equilibrium fraction of dormant cells in the tumour (in the absence of treatment) is *σ/*(*σ* + *ϱ*). A plot depicting this maximum fraction as a function of *ω* and *d*_*a*,daily_, is given by Figure 3.

**Figure 3.**
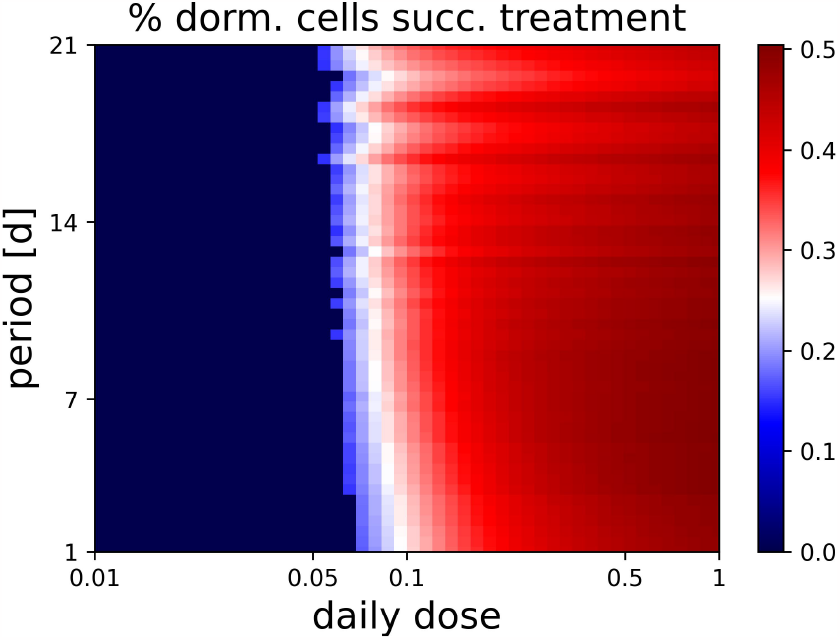
Maximum fraction of dormant tumour cells such that treatment is still successful (*J*_1_ *<* 100) for *ϱ* = 0.05 and *σ*_*T*_ = 0, with *ω ∈* [1, 21] and *d*_*a*,daily_ *∈* [0.01, 1]

Quite strikingly, we see that tumours with more than a fraction of 0.5% dormant cells cannot be treated effectively with a single drug under the imposed constraints on the maximum daily dose. The situation gets even worse considering the possibility of drug-induced switching from active to dormant as a result of stress caused to the cells by the drug (*σ*_*T*_ *>* 0). This allows us to formulate a first general observation (GO):

> *Even a small fraction of cancer cells exhibiting short-term dormancy can prevent treatment success*. (GO 1)

### 3.3. Optimizing single-drug treatment protocols in the face of dormancy

Even though a single drug targeting active cells only may not be able to eradicate the tumour / cancer cell population entirely, it is still interesting to see how far one can optimize the treatment regime with respect to the above constraints (3.4).

We begin by optimizing the *tumour size at the end of therapy*. To investigate the effect of the treatment scheme, we plot the objective function *J*_1_ from (3.2) as a function of the constraints. This is done in the following figure with parameter values as before.

The most prominent feature in Figure 4 is the lack of treatment success. No allowed daily dose and no possible period can reduce the average number of cells over the last 30 days of treatment to below 100. In view of Figure 3 this comes as no surprise because through spontaneous dormancy we initially have 1*/*6 of the tumour in a dormant state. As one would expect, a large daily dose is associated with better treatment outcome. However, note that *J*_1_ is not very sensitive to the choice of treatment period. While the minimum is at a daily dose of 1 and a period of approximately 3 days, there is a very large area in which the outcome of treatment yields a similar result with there being close to no difference between daily doses of 0.1 and 1. This is due to the finite exit rate of cells from the dormant population. In fact, since we start with 1.8 *·* 10^7^ dormant cells, the number of cells in a dormant state is bounded from below by

**Figure 4.**
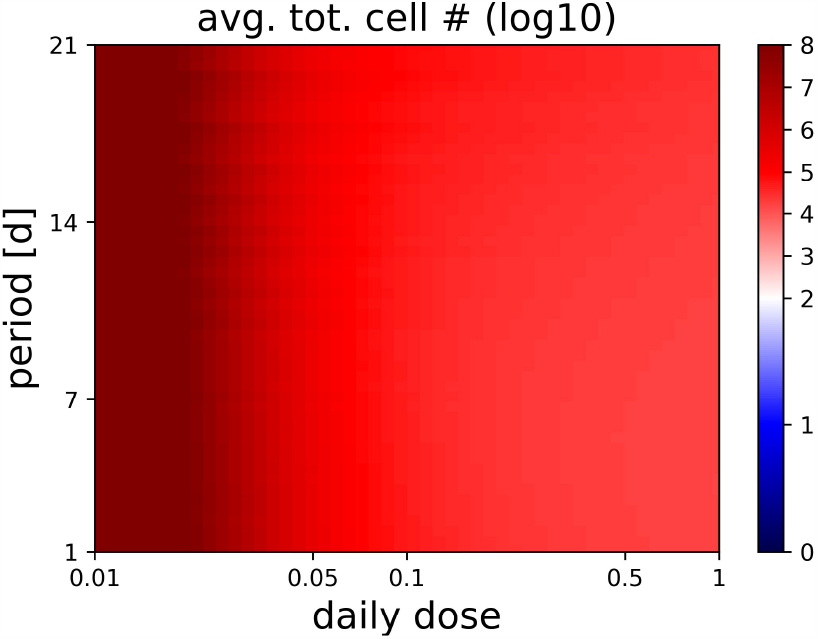
Heat map for the objective function *J*_1_ with spontaneous dormancy at rates *σ* = 0.01, *ρ* = 0.05 and drug-induced dormancy *σ*_*T*_ = 0.1.

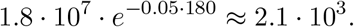

Interestingly, the optimal period is monotonically decreasing as the daily dose increases which we discuss in more detail in the appendix (Figure 13).

Turning to the *probability of the emergence of a resistance mutation* corresponding to objective function *J*_2_, there are three distinct mutation scenarios that one needs to consider:

- Mutations may only arise from active cells,
- mutations arise equally frequently from active and dormant cells,
- and the case in which mutations are more frequent in dormant cells.

The last case might appear somewhat non-natural, but [R+22] argue that that mutations in dormant cancer cells may actually occur at increased rates compared to active cells.

We will refer to these three scenarios as requiring a minimisation of the *active*, the *total* or a *weighted* area under the curve. For our basic study, we consider the cases *m*_*a*_ = 10^*−*8^ and *m*_*d*_ = 0, *m*_*a*_ = 10^*−*8^ and *m*_*d*_ = 10^*−*8^, and *m*_*a*_ = 10^*−*8^ and *m*_*d*_ = 10 10^*−*8^. Mutation rates of order similar to the reciprocal of the tumour size were used previously in [BCM+16] and rates of the order 10^*−*8^ appear in [G02] (when combining the rates of the two intermediate mutation steps). These choices result in the outcomes for *J*_2_ shown in Figure 5.

**Figure 5.**
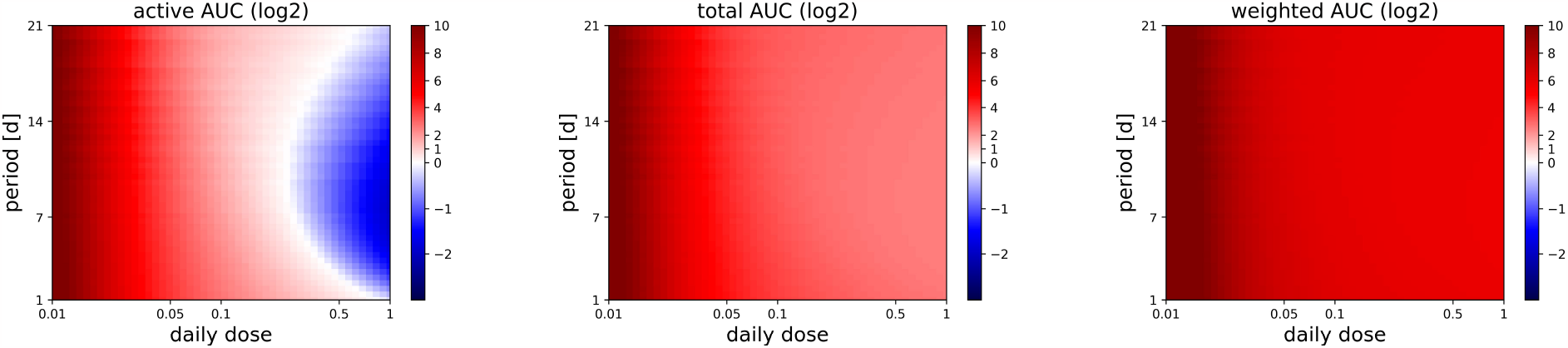
Values of *J*_2_ for the mutation probabilities *m*_*a*_ = 10^*−*8^ and *m*_*d*_ = 0 (left), *m*_*a*_ = 10^*−*8^ and *m*_*d*_ = 10^*−*8^ (middle), *m*_*a*_ = 10^*−*8^ and *m*_*d*_ = 10 *·* 10^*−*8^ (right).

Interestingly, as long as mutations can only arise from active cells, there is a region in which less than one resistance mutation can be expected to appear throughout the entire treatment. Also note that it seems most beneficial in this case to choose an intermediate treatment period of roughly 7 days. In particular, for a given daily dose, the integral of the active number of cells *J*_2_ is far more variable than the number of cells at the end of treatment given by *J*_1_, and highly dependent on the time interval between administrations.

In fact, for the active area under the curve we have differences which may be as large as a factor of However, if mutations may also arise from dormant cells, then we observe again a fairly indifferent behaviour of *J*_2_ to the treatment period. This is due to the almost linear decrease in the number of dormant cells as shown in the introductory examples from Figure 2. Hence, the treatment with a drug attacking only active cells is ineffective in this case.

To understand the variability of the active area under the curve, one may consider different treatment plans which result in the same final tumour size *J*_1_ but different AUC given by *J*_2_. An example for this is provided in Figure 6. There, both plans result in an average tumour size of roughly 2 *·* 10^4^ cells over the last 30 days of treatment. However, the daily treatment yields a total integral of the active number of cells of approximately 2 *·* 10^8^, whereas the treatment with a higher daily dose and a treatment interval of *ω* = 13 yields an integral of approximately 0.49 *·* 10^8^. This difference is due to the slow initial decrease in the number of active cells in the case of daily treatment. Since the initial large tumour size contributes the majority of the area under the curve, it is advantageous to reduce the tumour size quickly with a high initial dose. We may thus formulate our second general observation:

**Figure 6.**
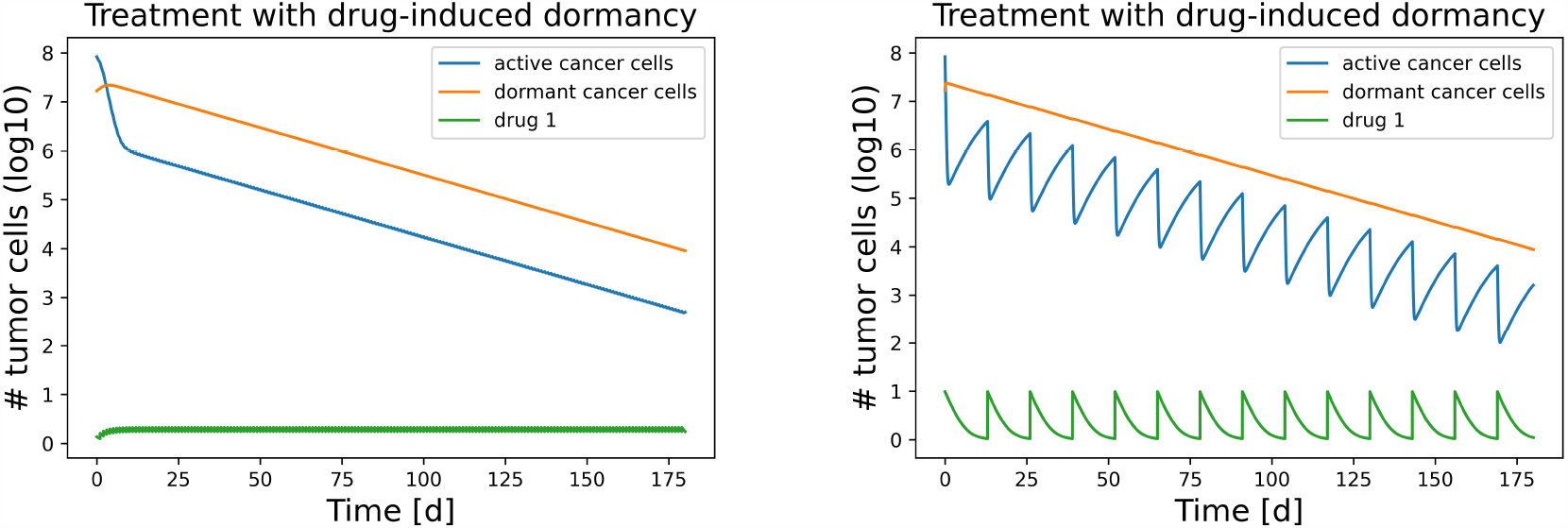
The development of the number of tumour cells over time with drug-induced dormancy at rate *σ*_*T*_ = 0.1. Left: *d*_*a*,daily_ = 0.37, *ω* = 1. Right: *d*_*a*,daily_ = 0.64, *ω* = 13.

> *Treatments with a single drug targeting only active cells can yield significantly different active AUC values J*_2_ *despite similar final tumour sizes J*_1_. (GO 2)

### 3.4. Combination treatments minimizing cancer populations

As we saw before, even under an optimized treatment plan with a single drug targeting active cells, the presence of dormancy may prevent successful treatment in the sense of achieving *J*_1_ *≤* 100. Therefore, it seems natural to additionally apply a second drug which directly affects dormant cancer cells. In [RM19, SD21] the suggested strategies to counter dormant cancer cells encompass three distinct strategies:

- increasing resuscitation rates by application of a “wake-up” drug,
- directly attacking the dormant cells,
- or artificially prolonging dormancy times.

We now discuss the effects of each of the three strategies separately.

#### 3.4.1. Increasing resuscitation rates

The wake-up strategy to reactivate dormant cells seeks to make dormant cells vulnerable to the first drug targeting active cells. In particular, this strategy may also counter adverse effects of drug 1 (which can also act as a trigger for dormancy initiation) and hence should increase treatment efficiency. However, the presence of a secondary drug increases the complexity of the associated optimal control problem. For example, there may be an advantage of administering the two drugs not at the same times but instead to incorporate a slight offset. To account for this, we introduce a parameter *t*_off_ *≥* 0 which determines the delay at which we administer drug 2 (compared to the administration of drug 1). For the optimal control problem this now gives a further constraint

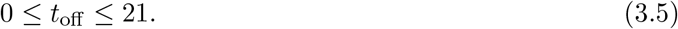

Note that we could in principle also ask about offsets for drug 1, but due to periodicity the areas in which an offset of drug 1 is useful will coincide with the areas which offset drug 2 by almost an entire period. Similarly, for shorter treatment periods, we will see the largest benefit along the lines *y* = *x/k* for *k ∈* ℕ and in the areas between these lines treatment outcome will be worse. This implies in particular that offsets *t*_off_ larger than *ω* will result in a treatment outcome which is similar to that with offset *t*_off_ mod *ω*. The only qualitative difference in these outcomes is the reduced effect of the second drug because it has not been used for several administration periods.

Regarding the objective *tumour size at the end of the treatment*, Figure 7 (left) quite strikingly shows that adding a wake-up drug can provide a very effective treatment scheme. Besides this basic observation, one can further see that incorporating offsets does not significantly improve the treatment outcome with respect to the final tumour size. This seems to be related to our modelling idealization that the administered drugs are immediately in the system with their maximum potential. Involving pharmacokinetics could change this observation. For shorter treatment periods, there are no large repercussions deviating from the ideal treatment for which both drugs are administered simultaneously. Moreover, for all periods and all offsets, the treatment outcome with respect to the objective *J*_1_ is consistently better than treatment with only a single drug. Only in a small area, this combination treatment is not successful by our definition of *J*_1_ *<* 100. This is due to the large offset where dormant cells are activated at a time when drug 1 is not effective anymore. However, even in this case combined treatment is still better than the single-drug scheme.

**Figure 7.**
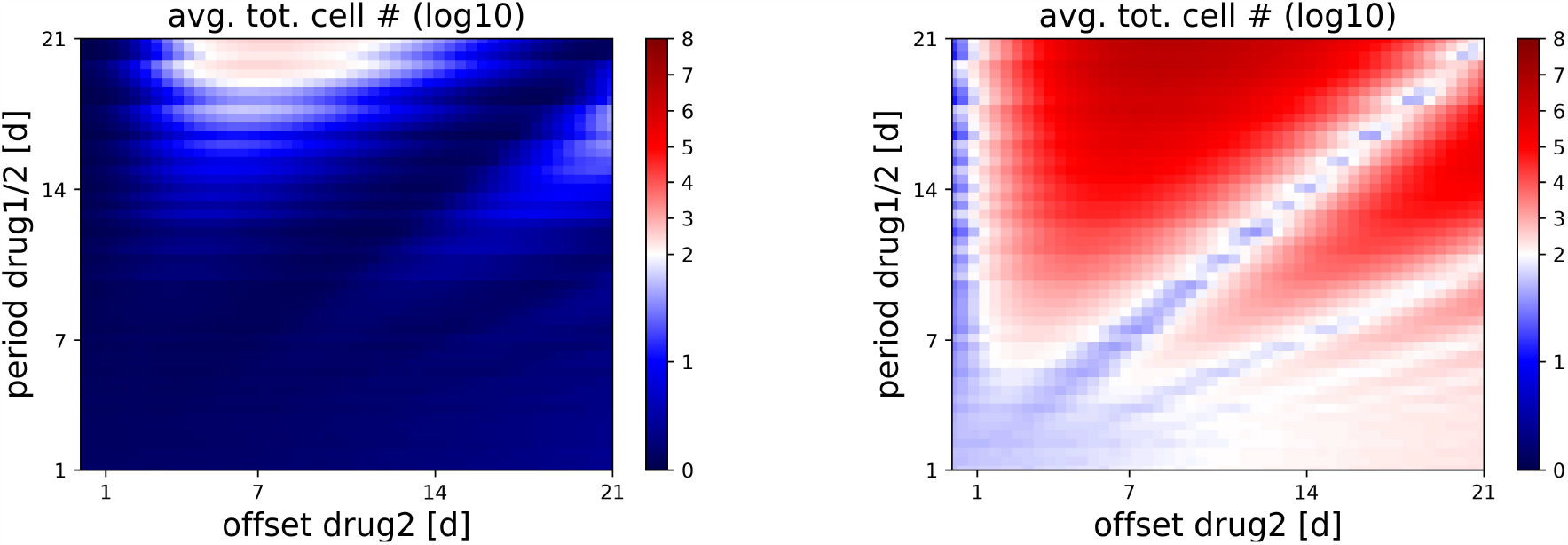
Heat map for the objective function *J*_1_ with combination treatment where the second drug activates dormant cells. Both drugs have a daily dose of 0.1. Left: *σ*_*T*_ = 0.1, *ϱ*_*T*_ = 0.5. Right: *σ*_*T*_ = 1, *ϱ*_*T*_ = 1.

If the drug-induced switching rates *σ*_*T*_ and *ϱ*_*T*_ are much larger though and hence switching is induced faster, the picture changes in an interesting way. Here, it becomes imperative to administer the drugs simultaneously as seen in Figure 7 (right). Observe that despite the second drug merely countering the induced switching of the first drug, there are again areas in which the tumour can be treated successfully. This is surprising, considering that the presence of spontaneous dormancy alone was preventing successful treatment. In particular, this indicates some synergistic effects between the two drugs. Observe though that there are now also areas in which the treatment outcome with two drugs is *worse* than with a single-drug treatment, for the same reason as in the left image. This leads us to formulate the following two general observations:

> *A combination treatment with a “wake-up” drug acting on dormant cells can be highly succesful regarding the objective function J*_1_ *even in scenarios where single-drug treatment clearly fails*. (GO 3)

> *For combination treatment with a “wake-up” drug acting on dormant cells, and high induced switching rates it seems best to administer both drugs simultaneously, i.e. with no time-offset*. (GO 3’)

We did not exhaust the constraint (3.4) on the maximum daily dose by far and already saw a significant improvement to what we call treatment success when the drugs were administered simultaneously. Increasing the drug doses then begs the question how important the mixture of the drugs is to obtain good results. We visualize this in Figure 8 (left). Here, we see that the area which eradicates the tumour encompasses almost all mixtures of the two drugs with a total daily dose of 1 except for those which consist almost exclusively of one of the drugs. In particular, the treatment is again not sensitive to changes in the period.

**Figure 8.**
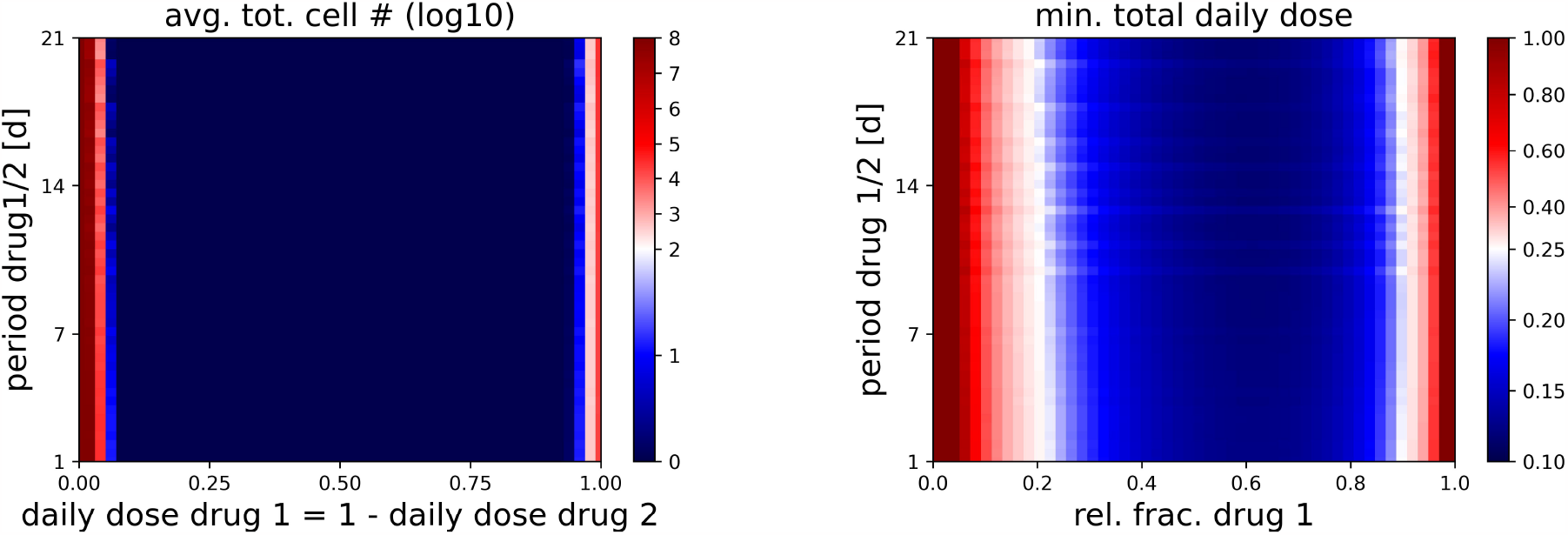
The objective *J*_1_ for various periods and combinations of daily doses for both drugs being administered simultaneously with the resuscitation approach for *σ*_*T*_ = 0.1, *ρ*_*T*_ = 0.5. Left: Value of *J*_1_ for given treatment plan. Right: Minimal total daily dose for given ratio of drugs 1/2 for *J*_1_ *≤* 100.

While we see that the maximal allowed total daily dose of 1 leads to a successful treatment outcome for many of the possible combinations, it may also be of interest to determine the minimal total dosage for a given mixture of the two drugs which leads to successful treatment in order to minimise negative side effects. We show this in Figure 8 (right), where we draw the boundary of the divergent colour scheme arbitrarily at 0.25 which corresponds to only requiring a quarter of the maximum allowed daily dose. Because drug 2 does not contribute to the elimination of cells directly, it is natural to find the minimum required total dose for treatment success to be for a mixture with slightly higher concentration of drug 1 than drug 2. We also observe that towards the boundaries of the drug mixture space, i.e. for those drug administrations which consist of almost exclusively one drug, the decline in the minimum total daily dose is fairly steep. This is indicative of the synergistic effects of the two drugs.

Concerning the *emergence of a resistance mutation* however, we find that the area under the curve depending on the mutation rates does require more precise treatment plans for success. We show treatment success for the three mutation scenarios from the previous subsection in Figure 9.

**Figure 9.**
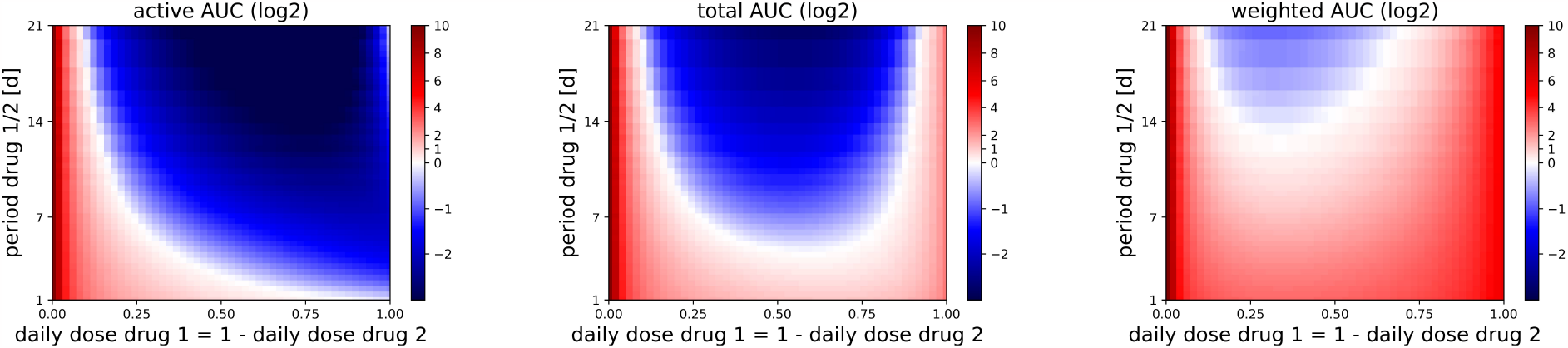
Values of *J*_2_ for the mutation probabilities *m*_*a*_ = 10^*−*8^ and *m*_*d*_ = 0 (left), *m*_*a*_ = 10^*−*8^ and *m*_*d*_ = 10^*−*8^ (middle), *m*_*a*_ = 10^*−*8^ and *m*_*d*_ = 10 *·* 10^*−*8^ (right) with simultaneous administration of both drugs and resuscitation rate *σ*_*T*_ = 0.1.

We see that longer treatment periods *ω* are consistently favourable to minimize the expected number of resistance mutations, regardless of the relative weighting of the two drugs. In particular, when dormant cells are able to mutate, a sufficiently long treatment period is mandatory to reduce the chance of emerging mutations below our threshold *J*_2_ *<* 1. This is due to the nature of the interplay of daily doses and periods: For the same daily dose, a longer period implies a higher administered dose at the beginning of each cycle. This in turn leads to a more rapid decline immediately after administration. However, due to drug decay there is also a longer period of potential recovery for the tumour. But during this period, the tumour is of significantly smaller size than at the time of drug administration, and hence, the recovery phase has a relatively minor impact on the area under the curve. We obtain another general observation:

> *To minimise the probability of an emerging resistance mutation, it is advisable to administer large doses rarely as opposed to administering small doses frequently*. (GO 4)

#### 3.4.2. Directly attacking dormant cells

So far we have only been dealing with dormant cells indirectly by reactivating and making them susceptible to the drug targeting active cells. However, there is also evidence that some drugs may be able to kill dormant cells directly [RM19]. We assume the killing rate for dormant cells to be lower than the killing rate for active cells, and also to be lower than the induced switching rate. To reflect this, we choose the drug-induced death rate of dormant cells to be *μ*_*dT*_ = 0.1.

Regarding the *final tumour size* at the end of the treatment, the objective function *J*_1_ subject to (3.4) takes the values shown in Figure 10.

**Figure 10.**
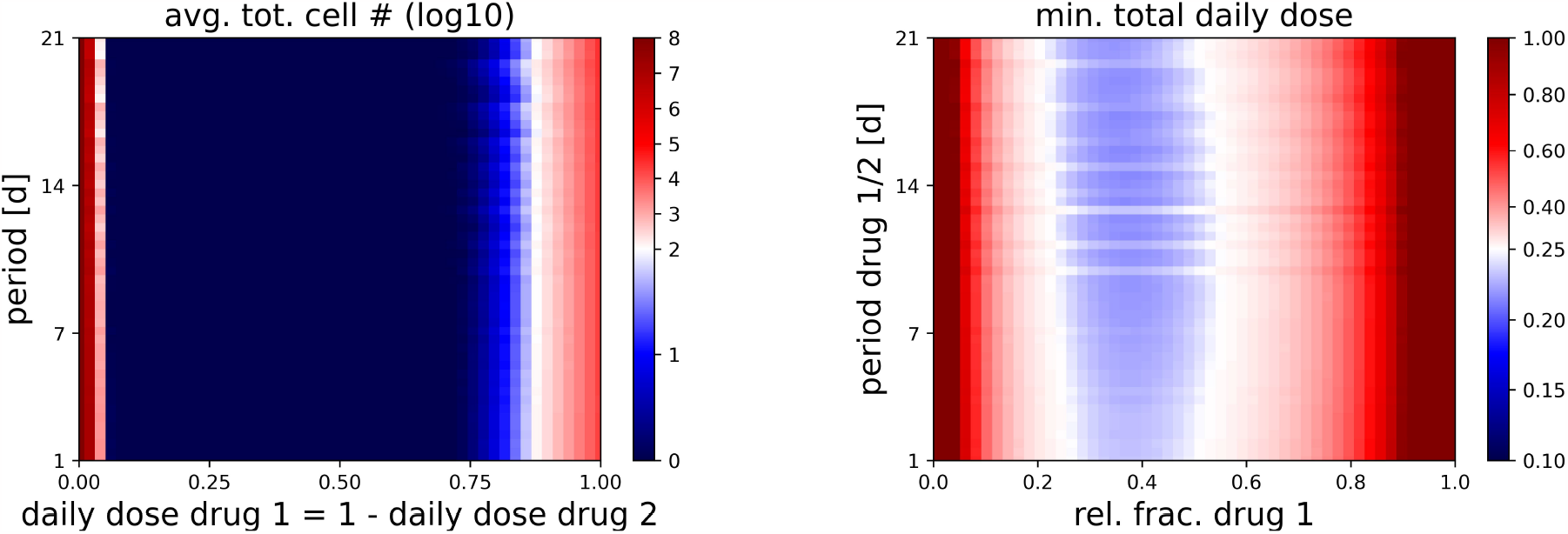
The values of the objective function *J*_1_ for different periods and convex combinations of daily doses in the strategy targeting dormant cells with rates *σ*_*T*_ = 0.1 and *μ*_*dT*_ = 0.1. Left: Value of *J*_1_ for given treatment plan. Right: Minimal total daily dose for given ratio of drugs 1/2 for *J*_1_ *≤* 100.

Note that it is always best to make use of the maximum allowed daily dose for which we consider different distributions of the total dose into drug 1 and drug 2 in Figure 10 (left). As is easily seen, despite drug 1 and drug 2 both on their own being highly inefficient at diminishing the number of cancer cells, considering a suitable mixture of both eradicates the tumour entirely. In fact, this is true for a wide range of mixtures. Again, the treatment is fairly indifferent with respect to the period. When considering the minimal daily dose that is needed for successful treatment in the sense of *J*_1_ *≤* 100 we see that this dose, perhaps surprisingly, is much larger than compared to the case of the wake-up strategy (Figure 8) despite their similarities of the treatment outcome when considering the maximum total daily dose. We arrive at the following:

> *For the killing strategy, a range of mixtures of the drugs can lead to near complete tumour remission, despite each drug being inefficient when on its own*. (GO 5)
>
> *For both the wake-up strategy and the killing strategy, successful treatment can be possible at low doses if mixtures are chosen optimally. The wake-up strategy can be efficient at a lower dose than the killing strategy*. (GO 5’)

The observation regarding comparisons of different strategies is of course highly dependent on the exact parameter choices. We will look further into this in the discussion section 4.

While the behaviour of this treatment approach with respect to the objective *J*_1_ is reminiscent of the one from the wake-up strategy, with regard to the *J*_2_, there are some intriguing differences as shown in Figure 11.

**Figure 11.**
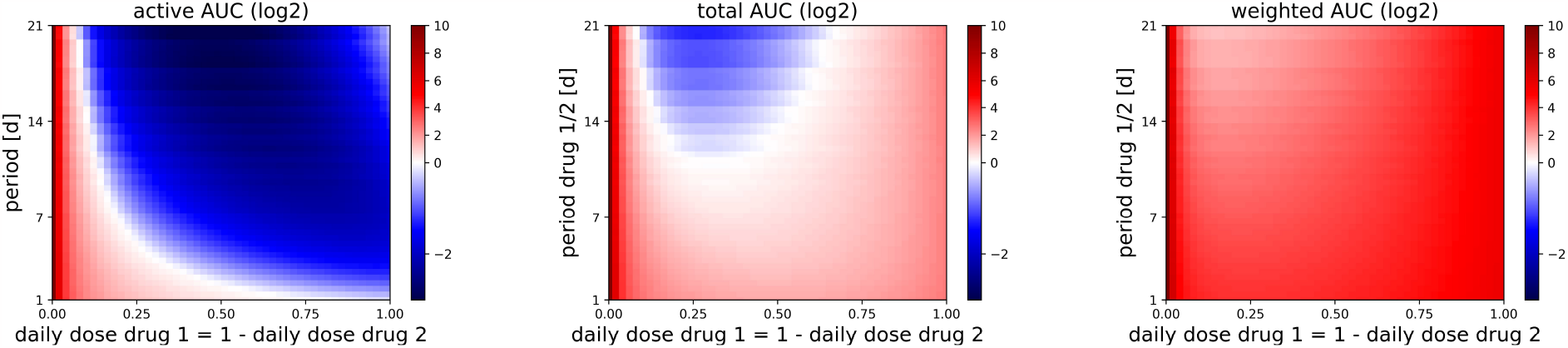
Values of *J*_2_ for the mutation probabilities *m*_*a*_ = 10^*−*8^ and *m*_*d*_ = 0 (left), *m*_*a*_ = 10^*−*8^ and *m*_*d*_ = 10^*−*8^ (middle), *m*_*a*_ = 10^*−*8^ and *m*_*d*_ = 10 *·* 10^*−*8^ (right) with simultaneous administration of both drugs for targeting dormant cells with rate *μ*_*dT*_ = 0.1.

As we can see, for our choices of parameters, the killing strategy is much less efficient in reducing the area under the curve of the dormant population when compared to the wake-up strategy, despite effectively eliminating the tumour. In the case where mutations are more frequent in dormant cells, no combination of drugs within our constraints is particularly efficient in preventing the emergence of mutations. However, when mutations are equally likely to occur in active and dormant cells, then there is a small area in which the emergence of mutations during treatment can be reduced significantly. The main observation is:

> *If mutations occur frequently in dormant cells, then in order to reduce J*_2_ *it seems better to employ the wake-up strategy instead of attacking the dormant cells directly*. (GO 6)

#### 3.4.3. Decreasing resuscitation rates

The idea of extending dormancy periods by decreasing the resuscitation rates of dormant cells requires an approach that is entirely different to the previous two schemes. Indeed, instead of aiming to decrease the size of the tumour, one wishes to suppress the tumour regrowth after an initial treatment targeting active cells. Here, we sketch this strategy by first applying treatment with drug 1 which targets only active cells to decrease the tumour size as much as possible. This happens for *T*_1_ = 180 days. Then, we apply the second drug, which is responsible for keeping the cells dormant for another extended period of time, say again *T*_2_ = 180 days. However, this alone is not enough to keep the tumour from regrowing since the initial active population may not have been completely diminished. We thus further need to assume that the second drug also encourages active cells to enter a dormant state, which seems somewhat justifiable for a drug which alters the environment in favour of sustaining dormancy. Note that it does not seem too prudent to keep cancer cells dormant at the beginning of treatment since this would prevent the tumour from entering remission in the first place, so that treatment by drug 2 will really only begin from time *T*_1_ onwards. This leads us to reconsider the system (2.3) and modify it to read

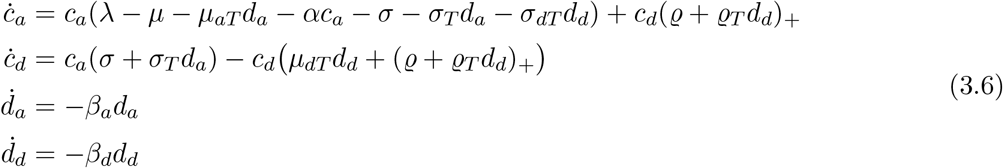

on the time interval [*T*_1_, *T*_1_ + *T*_2_], where *σ*_*dT*_ *>* 0. The convergence result of Lemma 2.1 then still applies on the entire interval [0, *T*_1_ +*T*_2_] by applying the lemma on the intervals [0, *T*_1_] and [*T*_1_, *T*_1_ +*T*_2_].

To analyse this approach, we assume a fixed treatment with drug 1 and vary treatment with the second drug. Again, we are interested in keeping the tumour as small as possible. However, while the objective *J*_1_ still measures this and could be used to make informed decisions about optimal use of drug 2, we will slightly modify it in order to allow for better comparison, independent of the success of drug 1. Suppose that treatment with drug 1 is performed for a time period *T*_1_ and treatment with drug 2 is then immediately continued for an additional time *T*_2_. Then we consider

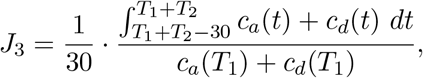

that is, we consider the relative increase of the number of cells averaged over the last 30 days of treatment with the dormancy drug compared to the number of cells present at the end of treatment with drug 1. This gives us a measure of how strongly drug 2 prohibits tumour growth. An example how the administration period and daily dose of drug 2 affect tumour regrowth is shown in Figure 12.

**Figure 12.**
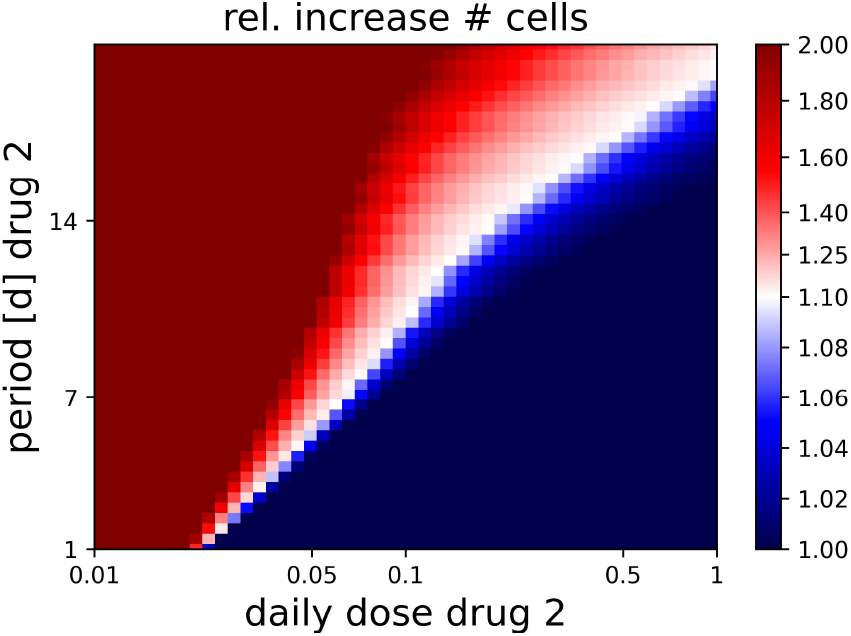
The values of the objective function *J*_3_ for different periods and doses. Parameters are *T*_1_ = *T*_2_ = 180 days, *ω*_1_ = 14 the treatment period for drug 1, daily dose *d*_*a*,daily_ = 1, *σ*_*T*_ = 0.1 for drug 1 and *σ*_*dT*_ = 1 for drug 2 and *ϱ*_*T*_ = *−*1.

We claim an increase by a factor of 1.1 over the period of *T*_2_ = 180 days as treatment success, since this yields a doubling time of about 4 years, which should be tolerable for most applications. This approach is heavily reliant on the first drug eliminating most of the tumour. In our example, at time *T*_1_ there are a total of roughly 7800 cells left. These then also play only a small role in the emergence of a resistance mutation compared to the 10^8^ cells that we considered at the beginning of treatment.

The figure shows that treatment can be successful even for relatively small doses of the dormancy-encouraging drug. As opposed to treatment with only one drug or the wake-up strategy with two drugs, it is optimal for all doses to have the minimal treatment period of *ω*_2_ = 1 days. This leads to our final general observation:

> *The strategy to inhibit tumour regrowth can be quite successful, and is best when treatment is applied frequently in small doses. It is already effective for relatively small daily doses*. (GO 7)

## 4. Discussion

In this section, we first review our modelling assumptions, then comment on the plausibility and robustness of our concrete parameter choices, followed by a discussion of our main results (in particular our ‘general observations’). We conclude with a short review of related models, and briefly sketch areas for possible future research.

### 4.1. Basic modelling assumptions

We provided a stochastic individual-based (toy) model for the dynamics of a population consisting of actively reproducing and dormant (non-proliferating) cancer cells. The set-up using a two-type birth-death process with interaction for reproduction and competition is standard in the literature. Our choice of dormancy mechanisms however is quite specific. Here, we assume “short-term dormancy”, by which we mean that dormancy initiation and resuscitation happen with rates comparable to reproduction and decay rates in the model (instead of other forms of dormancy mentioned in the introduction). Our dormancy transitions are thus reminiscent of *phenotypic switches* which are similarly (and much more commonly) modelled in the literature as we will discuss below in Section 4.4. Note however that dormancy has rather particular features, including protection from the environment, and usually complete absence of reproduction (i.e. binary fission and cell death).

Whether individual cell dormancy of such a kind really exists in cancer is of course question for empiricists – here, we take the freedom of theorists to merely postulate several paradigmatic mechanisms related to short-term dormancy, and investigate their hypothetical effects on the population dynamics of cancer cells, including their consequences for different treatment approaches and schemes.

Our model is thus certainly an idealization and in particular a rather drastic over-simplification of cancer cell population dynamics. However, there is some value in regarding and analysing highly reduced models, since they allow one to observe some of the fundamental effects of dormancy in a very clear fashion, without other possibly confounding factors. They also allow for rather general observations, independent of the concrete cancer type and the many other specifics in cancer treatment. If one observes interesting and paradigmatic behaviour already in simplified models, then this allows to hunt for similar effects in subsequent much more realistic and fine-tuned models.

We will discuss below some of the many possible extensions of the model to cover more specific scenarios, including further cell types, spatial interactions and structure, etc.

### 4.2. Parameter choices

We now turn our attention to the plausibility and robustness of our model parameters. Recall that at the beginning of Section 3.2 we fixed most parameters of our model and later also picked the drug administration period and drug dosage. While we are not trying to model a concrete type of cancer, we still want to ensure that these fixed parameters are somewhat reasonable.

- For the *initial number of cancer cells*, we choose 10^8^ cells. This roughly corresponds to the minimal clinically detectable tumour size [D09, E12, AK+20]. Alternatively, we can interpret this as a small residual tumour tissue after surgical removal. Note that, since we study small tumours and set the competition parameter to zero, the choice of the initial cancer cell number does not qualitatively change the system behaviour but merely scale the later number of cancer cells proportionally.
- Regarding the *cancer growth parameters λ* and *μ*, the net growth rates *λ − μ* of untreated tumours vary between 0.008–0.25 per day, while the cell loss factor *μ/λ* ranges between 0.69 and 0.93 [IIZB74, T89]. We choose *λ* = 0.5 and *μ* = 0.45 to lie within these ranges and represent a tumour type that is neither extremely aggressive nor extremely slow-growing.
- Concerning the *drug decay rates*, we chose both of the drugs to have an identical decay rate for the sake of simplicity. If the drugs have significantly different decay rates, then our observation regarding the offset between drug administration times may not hold true. For example, if the drug waking up dormant cells has very slow decay and hence remains at a high concentration for long periods of time, then it may be beneficial to administer this drug several days before the drug killing active cells is administered. Similarly, this may apply to the killing. Our choice of decay rate *β*_*a*_ = *β*_*b*_ = 0.4 does not represent a specific drug but lies within the range of typical parameters for small molecule and other biological drugs approved for oncology indications [LD17].
- With regards to *dormancy related spontaneous transitioning rates*, the literature seems to be scarce. Further, there appears to be no consistent definition of what exactly dormancy is. E.g. for cells it is difficult to distinguish between a resting period in the cell cycle (referred to as *quiescence*) and a ‘truly dormant state’. In order to account for these ambiguities, within the context of this paper, dormancy should refer to a prolonged period without reproduction in the absence of drugs that influence these dynamics. This leads us to choose the resuscitation rate to be relatively low at *ϱ* = 0.05, corresponding to an average time of 20 days spent in the dormant state. Further, we assume that dormant cells should not comprise the majority of a tumour and hence we settled on *σ* = 0.01, that is, any cell enters a dormant state on average after 100 days. It is very possible that such long dormancy periods are entered spontaneously even less frequently and hence one would need to choose *σ* smaller. Doing so would decrease the dormant population in the beginning of therapy and hence make any treatment plan attacking mostly active cells more effective.
- Similarly, there seems to be only very little literature regarding *induced switching rates*. There are studies which describe the exhibition of certain dormancy markers within several days of treatment [TJC+17, KS+22]. Again, it seems somewhat unclear what a precise definition of dormancy should be. We decided to quantify the effect of induced switching by the drug targeting active cells (usually) to be one tenth of the killing rate. This is reasonable if one believes the ‘stress induced’ switching to only occur as a side effect. For the strategy keeping cells dormant, the induced switching rate from the active to the dormant state is of larger magnitude since this is intended purpose of the drug. All of these drug-induced rates are, however, somewhat arbitrary, since they scale with dosage for which we do not have any realistic assumptions. For the drug targeting dormant cells, we assumed that killing dormant cells is more difficult than killing active cells and hence chose the corresponding death rate at 1/10 of that for active cells.

Given the uncertainties related in particular to the choice of the dormancy-related parameters and drug dosages, this begs the question how robust our results are given these unknowns. We can confidently say that treatment with only one drug will perform better the less induced switching it causes. However, as shown in Figure 3, the fraction of dormant cells must be vanishingly small in order to see successful treatment. Hence, we believe that our model will still show dormancy to be a factor preventing successful treatment for a large range of parameters.

Our observations regarding the area under the curve (i.e. total number of cells throughout treatment) mandating large doses and long treatment intervals for combination therapy will hold true regardless of the exact rates of the drugs. It may be the case though that under different parameters it is not possible to obtain any admissible treatment plan for which the expected number of mutations is sufficiently low.

For combination treatment with a secondary drug targeting dormant cells, we believe the second drug to still yield a major improvement for treatment outcome when considering the wake-up or killing strategy. Of course it is difficult to incorporate confounding factors of the drugs interacting with each other into our model as well as costs associated with the drugs from e.g. side effects. As such, it is questionable what limits there are regarding the total dose that can be administered at each treatment time and what the total area under the curve of the drugs is allowed to be. Such trade-offs could heavily influence the outcome of the optimal strategy. However, as pointed out by [KCV21], it is more important to have a large area of parameters which are similar to the optimal strategy as opposed to exactly pinpointing the optimum, particularly because we aim to find rules of thumb from our analysis.

We also found that for the parameters we chose, one often does not necessarily need to apply the maximum allowed daily dose for successful treatment. As discussed in [MBSX22], this opens up the possibility of treatment schedules other than the frequently used maximally tolerated doses.

In conclusion, we believe most of our findings from this model to hold true in a qualitative sense even under variable model parameters. However, the extent to which combination treatment can improve therapy success may be rather sensitive to the parameter choices.

### 4.3. Main results

We now discuss some aspects of our main results, in particular our ‘general observations’ from Section 3.

#### The role of short-term dormancy in treatment failure

Our first main point is that our model demonstrates the importance of spontaneous short-term dormancy in cancer treatment already in our idealized theoretical setup. As stated in (GO 1), classical single-drug therapy consistently fails when small fractions of the tumour are in a dormant state. This holds both for the objective ‘minimal tumour size at end of treatment’ as well as ‘suppression of a resistance mutation’, although in the latter case, if mutations are absent in dormant individuals, there is at least some room for treatment optimization.

While it is *in practice* highly unlikely that chemotherapy on its own has any chance to eradicate a tumour, our idealised scenario shows that spontaneous as well as treatment-induced dormancy should generally worsen treatment outcomes significantly. However, in reality there will likely be further factors leading to and impacting on the presence of dormancy, whether that is hypoxia or lack of nutrients or factors from the micro-environment or cell-cell interactions [AG07, DRS21]. While there is a term involved in our model which would be capable of capturing competition for resources, this is a *global* term, whereas the environment impacts the cells locally. It would require a spatial model to incorporate such environmental factors.

#### Initial dosages and the chances of emergence of resistance mutations

A second major takeaway is concerned with the emergence of a resistance mutation during treatment. For simplicity we have assumed the rate of mutation to be constant throughout treatment. However, there also seems to be evidence that the presence of treatment may elevate mutation rates in dormant cancer cells [R+22], which is an observation that we incorporated in a simplified form by having an overall increased weighting on the dormant area under the curve. While we do not know for certain how fast mutations arise in cancer cells, what we can say independently of this is that in order to reduce the risk of the emergence of a mutation in cancer cells, it seems generally advisable to diminish the number of present cells as quickly as possible. Since the number of cells is largest at the beginning of treatment, this typically demands an initial dose that is as large as possible.

Because our somewhat restrictive modelling approach assumes the same dose for each treatment cycle and the integral of the drug concentration to remain constant for variable treatment cycle length, this often results in large doses given at large intervals, as we noted in particular in (GO 4) for the wake-up strategy. The same observation also applies for directly eliminating dormant cells. Interestingly, for single-drug therapy, as shown in Figure 5, as long as mutations can only arise from active cells it is ideal for the maximum dosage to have a treatment period of roughly 7 days as opposed to the longest or shortest admissible period.

#### Single-drug treatment may yield different results regarding the objectives J_1_ and J_2_

This result is related to (GO 2). Usually, a drug will at first force the tumour to shrink but after some time the drug has decayed enough that the tumour can grow again. In order to minimize the probability of a resistance mutation, it is not too important how much the tumour size declines between drug administrations, but it matters how fast the tumour is diminished in the beginning while the drug concentration is still high. Therefore, we may have treatment protocols which are efficient at reducing the probability for a mutation but have relatively large final tumour size. A particularly illustrative example for this is shown in Figure 6 (right), where we see that treatment exclusively with a single drug and a mutation probability *m*_*a*_ = 10^*−*8^ sees on average less than one mutation from active cells but the final tumour size is of the order 10^4^ cells since the majority of cells are in a dormant state. As soon as we take into account dormant cells however, it is only possible to get small mutation probabilities when also the final tumour size is small.

#### Combination treatments: Simultaneous administration vs offset between different drugs

Our next point is that combination treatment in our model is most effective when the drug targeting active cells and the drug targeting dormant cells are administered simultaneously and when done so, a wide mixture of such treatment plans yields desirable outcomes. We commented on this in (GO 3’) and (GO 5). This seems to be to a large degree a result of the instantaneous effectiveness of our drugs. Despite the optimum being a simultaneous administration, we showed in Figure 7 that there is some tolerance when considering a delayed administration of one of the drugs which still yields a successful treatment outcome. Such delays might need to be considered when the drugs are not fully compatible with each other and may result in undesirable side effects if administered at the same time or when different pharmocokinetics result in the drugs being distributed in the body at different speed, i.e. it would be practical to administer them simultaneously but have them spread at slightly different speed.

#### Targeting dormant cells: Wake-up vs. killing strategy

A further point is that regarding treatment outcomes, the wake-up strategy and the killing strategy are very similar. The reason for this is that both approaches essentially increase the death rate of dormant cells. However, we found a subtle difference with (GO 6) concerning the emergence of a mutation. If mutations are more frequent in dormant cells, then it is advisable to resuscitate the dormant cells instead of trying to kill them directly. Of course, this is an observation which is highly sensitive to the parameter choices that we discussed above. In our example, the effect is due to the fact that we assumed inducing a switch to the active state to be more easily accomplished than eliminating a dormant cell directly.

#### Targeting dormant cells: Extending dormancy periods

Our last point concerns the strategy to keep cancer cells dormant. This approach can realistically only be considered when prior treatment has been able to reduce the size of the tumour to an acceptable level. The dormancy strategy is then aimed at prohibiting tumour regrowth and hence may be useful when further treatment with more aggressive drugs is not feasible, e.g. due to undesirable side effects. In such a scenario, our model predicts that it is best to keep the drug at a low constant level (realised by administering the doses daily). Such a treatment strategy then inhibits tumour growth fairly effectively for small dosages as described in (GO 7).

### 4.4. Comparison to other models

There is a near endless number of suggested models for cancer growth in idealised scenarios [YMvH+19] as well as models for treatment of cancer [EBE11, KCV21] and other applications such as the emergence of resistance mutations and metastasis in e.g. [ALM15] and references therein. Our model is motivated from stochastic interactions of individual cells, but the approximation of the dynamics from Lemma 2.1 shows that we are in a setting with classic logistic growth. Also, the models for treatment usually are *concentration-dependent* meaning that the dynamics are depending on the concentration of the drug and cancer cells, which is the case that we adopted into our setting as well.

While there is an abundance of models for cancer growth, only few of them take dormancy at the cellular level into account. In most of the models contained in the references above, the term dormancy refers to *tumour mass dormancy* as opposed to cellular dormancy. Explicit models of cellular dormancy in cancer were discussed for example in [PU05, KW07, SRC+19] and references therein. However, these are not concerned with identifying optimal treatment plans. Instead, they are more focused on equilibria and time to extinction under constant treatment. One of the closest models to ours discussed in the literature seems to be [G02], where dormancy in the form of quiescence and multi-drug chemotherapy is considered but without specifically targeting dormant cells. The recent evidence for the possibility of directly targeting dormant cancer cells in various ways [RM19, SD21] is not included in any of those models.

More prominent is the case of phenotypic switching as discussed in [GV03, BCM+16, GK+19, GDLF20, AGKS22] and references therein. Such a model is from a mathematical perspective similar to that of dormancy, since dormancy is a phenotype of a cell which does not proliferate and may be resistant to specific chemotherapeutics. There are however a number of differences, since we for example assume dormant cells to not reproduce and also to not die unless targeted by a drug. Such an assumption is crucial for the treatment strategy which extends the dormancy times to be viable. In addition, a systematic study of the different strategies to counter dormant phenotypes seems to be missing in the literature also in the general context of phenotypic switching.

### 4.5. Future model extensions and outlook

An additional layer of complexity in cancer population biology is represented by the presence of a multitude of cancer cell types. A basic distinction is the one between “ordinary” cancer cells (CCs) and cancer stem cells (CSCs), where the latter may share certain features of dormant cells (cf. the results of the mouse experiment of [Z14]). In our basic model, we may incorporate both cancer cells (CC) and cancer stem cells, and allow for a variety of specific dormancy related and reproductive mechanisms in both classes according to the above sketched typology. CCs and CSCs may also react in distinct ways to drug administration, depending also on their state of activity/dormancy. One of the intricacies of such a model is the fact that CSC may exhibit symmetric but also asymmetric cell division, that is, they may produce a stem cell or a non-stem cell as a result of cell division.

Typical assumptions are that long-lived cancer stem cells initiate a tumour and produce (differentiated) cancer cells. The latter may still proliferate (but typically with a more limited proliferative potential, see e.g. [W+14]), but normally cannot produce stem cells. This leads to modelling approaches using multi-compartment models, which means that non-stem cells can divide only a finite number of times and move into the next compartment upon finishing cell division. However, there seems to be some evidence for plasticity in the direction of cancer cells changing back into a stem-like state [C+11] spontaneously. In addition, as observed by [VP15] in the case of radiation treatment, it is possible that treatment may also induce such a transition of cancer cells into stem-like cells.

Therefore, extending our model to include cancer stem cells, we immediately introduce a number of new possible transitions. In particular, one may require additional drugs targeting active (or dormant) cancer stem cells in order to eradicate the tumour.

Another way to enrich our basic model is to account for pharmacokinetics. In order to keep the model as simple as possible, we assumed the drugs to take immediate effect at the time of administration. However, in reality there is a delay between the injection of a drug into the system and its arrival at the tumour. Depending on the physiological circumstances, these delays as well as the absorption rates of the drug in the body can vary significantly between different drugs.

An aspect of the model which we overlooked for the purpose of simplification as well is the possibility that a drug does not only decay naturally but also gets absorbed and degraded by its interaction with cancer cells. This would mean that besides the linear decay we would see a degradation term depending on the drug concentration and the number of present cancer cells. Together with the above mentioned pharmacokinetics would create more involved drug dynamics and hence result in more realistic descriptions of the results of a treatment plan.

Furthermore, we simplified our model by assuming the constraint reflecting potential negative side effects to only be depending on the total drug dose. This could be more refined by account for different levels of toxicity of different drugs. In addition, as pointed out by [KCV21], more refined ways such as considering a population of healthy cells which are also attacked by the drugs and may not decline below a certain threshold could be implemented to obtain more precise constraints on the minimisation problem.

Another simplification we made was to assume that the drug doses and administration intervals were constant throughout treatment. One could remove this assumption and instead consider treatment protocols in which the drugs may be administered with different dosages at different administration times as in [ALM15]. This would allow for more elaborate treatment plans which in turn may result in different interaction patterns of the different drugs.

Lastly, a way to extend our model is to include other forms of therapy such as radiation or immunotherapy. In real-world applications, multiple types of treatment are commonly used in order to not only eradicate the primary tumour but also to target disseminated tumour cells and prevent metastasis. For example, radiation therapy can be incorporated by letting the death of cancer cells (and potentially induced switching rates) to be time-dependent with two possible values for the phases during which radiation is applied and when it is not applied.

## 5. Appendix

### 5.1. Scaling limit under discontinuous treatment - Proof of Lemma 2.1

*Proof*. We prove by induction over *i ≥* 1 that,

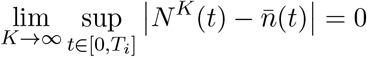

almost surely.

Consider the base case of *i* = 1. Then, the standard result of [EK86, Theorem 11.2.1] implies almost sure uniform convergence of *N* ^*K*^ to the solution of the ODE (2.3) 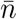 on [0, *T*_1_). At time *T*_1_, we have

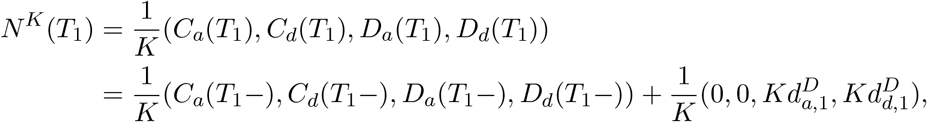

Since we have almost sure uniform convergence of *N* ^*K*^ on [0, *T*_1_), we find that

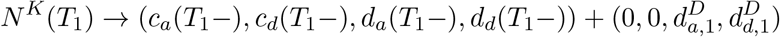

almost surely as *K → ∞*. The term on the right-hand side is exactly how we defined the solution 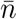 at time *T*_1_. Because 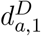 and 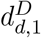are deterministic, the almost sure uniform convergence extends to [0, *T*_1_].

For the induction step we know by assumption that there is almost sure convergence of 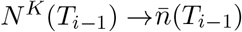. Hence we can again apply the standard Ethier–Kurtz result as in the base case to conclude our claim on [0, *T*_*i*_]. Because we have only a finite number of administrations, this finishes the proof. ?

### 5.2. Parameter choices

The following table lists the parameters values and ranges used to generate the simulations in this paper. For units, we measure time in days, cancer cells (active or dormant) in cell number *n*, and drugs (targeting active or dormant cancer cells) as a concentration *c*. Note that we do not prescribe a specific unit for the drug concentration (such as *μg/mL*) since we are not modelling a specific existing cancer drug and the concentration unit cancels against the drug effect rate units.

### 5.3. Additional illustrations

In Section 3.3 we pointed out that the optimal period between drug administrations for single-drug treatment is monotonically decreasing as the daily dose increases. We show this in Figure 13 Creating such a study has two confounding factors which impact the “optimal” interval between drug administrations. On one side, we need to discretise the space of admissible periods. In this case, we decided to do this by only allowing for administrations to occur at most every 0.5 days. On the other hand, the objective *J*_1_ averages over the last 30 days. This choice favours some periods over others since periods which have their last administration shortly before the end of treatment will have a lower average tumour size.

**Figure 13.**
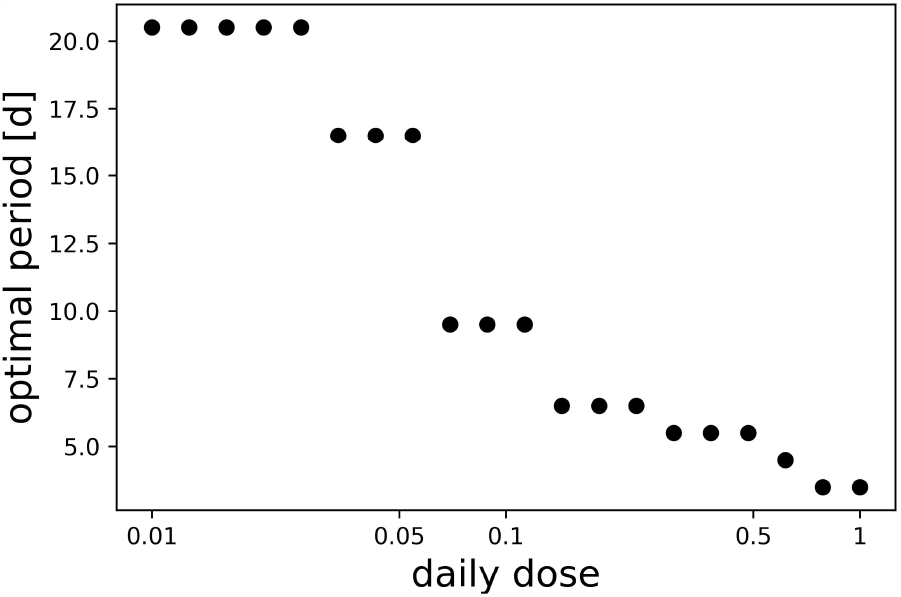
Optimal period for different daily doses which minimize the objective function J_1_ under treatment with a single drug.

## Acknowledgements

TP was supported by the Deutsche Forschungsgemeinschaft (DFG, German Research Foundation) under Germany’s Excellence Strategy *MATH+: The Berlin Mathematics Research Center*, EXC-2046/1, Project-ID 390685689. AT was partially supported by the ERC Consolidator Grant 772466 “NOISE”. All authors were partially supported by the Deutsche Forschungsgemeinschaft (DFG, German Research Foundation) under Germany’s Excellence Strategy *Haussdorf Center for Mathematics*, EXC-2047/1, Project-ID 390685813.

